# Performance of novel antibodies for lipoarabinomannan to develop diagnostic tests for *Mycobacterium tuberculosis*

**DOI:** 10.1101/2022.04.12.488063

**Authors:** Jason L. Cantera, Lorraine M. Lillis, Roger B. Peck, Emmanuel Moreau, James A. Schouten, Paul Davis, Paul K. Drain, Alfred Andama, Abraham Pinter, Masanori Kawasaki, Gunilla Källenius, Christopher Sundling, Karen M. Dobos, Danara Flores, Delphi Chatterjee, Eileen Murphy, Olivia R. Halas, David S. Boyle

**Affiliations:** PATH, Seattle, WA, USA; Global Health Labs, Bellevue, WA, USA; FIND, Geneva, Switzerland; Mologic, Thurleigh, Beds, United Kingdom; Departments of Global Health, Medicine, and Epidemiology, University of Washington, Seattle, WA, USA; College of Health Sciences, Makerere University, Kampala, Uganda; Public Health Research Institute Center, New Jersey Medical School, Rutgers University, NJ, USA; Otsuka Pharmaceutical Co., Ltd., Tokyo, Japan; Department of Medicine Solna, Division of Infectious Diseases, Center for Molecular Medicine, Karolinska Institutet and Department of Infectious Diseases, Karolinska University Hospital, Solna, Sweden; Mycobacteria Research Laboratories, Department of Microbiology, Immunology, and Pathology, Colorado State University, Fort Collins, CO, USA

## Abstract

Lipoarabinomannan (LAM), a component of the *Mycobacterium tuberculosis* (MTB) cell wall, is detectable in the urine of MTB infected patients with active tuberculosis (TB). LAM-specific antibodies (Igs) have been developed by a variety of traditional and recombinant methods for potential use in a rapid diagnostic test (RDT). We evaluated the analytical performance of the TB LAM Igs to identify pairs that offer superior performance over existing urine LAM tests. We assessed 25 new and 4 existing Igs in a matrixed format using a multiplex electrochemiluminescence-based liquid immunoassay. A total of 841 paired Ig combinations were challenged with *in vitro* cultured LAM (cLAM) derived from MTB strains representing diverse phylogenetic lineages, alongside urinary LAM (uLAM) from the urine of adults with active pulmonary TB. Analytical sensitivity of down-selected Ig pairs was determined using MTB Aoyama-B cLAM, while diagnostic accuracy was determined using clinical samples. When testing cLAM, the reactivity of Ig pairs was similar across MTB lineages 1-4 but lineage 5:6 had significantly more reactivity among Ig pairs. Overall, 41 Ig pairs had a strong binding affinity to cLAM, as compared to the reference pair of S4-20/A194-01, and 28 Ig pairs therein exhibited a strong affinity for both cLAM and uLAM. Retrospective testing on clinical urine specimens demonstrated varying sensitivities (12-80%) and specificities (14-100%). The five top pairs had a similar analytical limit of detection to the reference pair but in four instances, the sensitivity and specificity with clinical uLAM samples was poor. Overall, epitopes presented by uLAM are different from cLAM, which may affect antibody performance when testing uLAM in patient samples. Several new Ig pairs had similar ranges of high sensitivity to cLAM but overall, there were no new candidate Ig pairs identified in this round of screening with increased performance with uLAM as compared to an existing optimal pair.

## Introduction

Active tuberculosis (TB) is a leading cause of death from an infectious agent, despite being treatable with antibiotics (1). TB disease is caused by *Mycobacterium tuberculosis* (MTB). The distribution of TB is concentrated in low- and middle-income countries. In 2019, TB infected an estimated 10.0 million people globally, with an estimated 1.5 million deaths from TB (1). The COVID-19 pandemic has resulted in significant setbacks to reducing the global TB burden and in 2020 numbers of cases of TB diagnosed and reported fell to ~5.8 million cases (1). The mortality from TB has increased in 2020 and so providing access and provision for TB diagnosis and treatment is a global priority. Early diagnosis of TB and rapid treatment initiation are key for improved patient care and in reducing community transmission of disease (2). Many of the current diagnostic methods for TB infection are not ideal for rapid diagnosis. Collecting sputum can be challenging, poses risks to health care workers, and ineffective for diagnosing extra-pulmonary TB. Bacterial culture is most sensitive but can be slow and while smear microscopy is simple, rapid, and low cost, it has poor sensitivity (3). Molecular methods offer speed, diagnostic accuracy and may genotype resistance to some drugs but with higher costs for equipment, maintenance and being laboratory-based (4,5).

To improve the diagnosis of TB in high burden countries there is an urgent need for an effective non-sputum-based rapid diagnostic test or a triage tool that has the necessary performance to rapidly identify diseased patients for confirmatory testing and treatment (6–9). Lipoarabinomannan (LAM) is a lipoglycan that is a major structural component of the mycobacterial cell wall and also acts as an immunomodulator that may play a role in its pathogenesis (10,11). The structure of LAM from several mycobacterial species including MTB has been elucidated and mapped to 4 distinct domains (12–16). Soluble LAM is actively secreted from MTB bacteria and infected macrophages and is ultimately excreted via urine, an easy to access and relatively copious sample volume readily available from all suspected cases of TB (17–20). These factors make urinary LAM (uLAM) a candidate biomarker for a rapid antigen TB test (21). The terminal end of LAM has structural domains that present immunologic epitopes including at least one TB specific target (12,22,23), and uLAM is present in all active TB cases regardless of the anatomical site(s) of disease.

Recent advances to increase the sensitivity of LAM immunoassays have focused on developing Igs with improved sensitivity and specificity to TB LAM (22–26), sample preparation including pre-treatment to remove confounders (27–29), pre-enrichment of uLAM from urine prior to testing (30,31), amplifying the stripe signal (32) or highly sensitive immunoassay platforms (33–35). Our aim was to assess the analytical performance of TB LAM antibodies (Igs) to identify pairs that offer superior performance over existing urine LAM tests. We applied the MesoScale Diagnostic (MSD) immunoassay platform to interrogate 25 new Igs targeting LAM epitopes in both capture and detector positions in comparison to reference Igs previously screened on this platform (33). The optimal candidates were down selected based on their initial performance with cultured LAM from six different lineages of MTB and clinical uLAM, and the best in class then assessed for their limit of detection and performance with a test panel of clinical urine samples collected from South Africa, Peru, Uganda and Vietnam.

## Materials and Methods

### Antibodies and Antigens

A total of 29 Igs, including 4 previously referenced and 25 not previously screened, were assessed in this study. The Igs or their antigen-binding fragment (Fab) derivatives that target TB LAM were obtained from the Foundation for Innovative New Diagnostics (FIND), the Karolinska Institutet, Mologic, Otsuka Pharmaceutical, and Rutgers University (Table 1). These Igs included monoclonal forms in addition to recombinant Igs derived from phage display and recombinant forms manipulated from light and heavy chain sequences from other candidate anti-LAM Igs to create new Igs and Fab forms of Igs. These materials were generated from a variety of TB LAM antigen sources including purified LAM from *in vitro* cultured MTB cells, inactivated *in vitro* cultured MTB cells, synthetic glycans to specific TB LAM epitopes and the creation of recombinant forms prepared from memory B cells collected from convalescent TB patients (22). The purified *in vitro* cultured LAM (cLAM) used in this study included Aoyama-B LAM (Nacalai Tesque, CA, USA), H37Rv LAM (Biodefense and Emerging Infections Research Resources Repository Resources [BEI], Manassas, VA, USA), and from five TB strains representing lineages 1, 2, 3, 4 and 5:6 obtained from Colorado State University (See Dataverse file) (36).

**Table 1.**
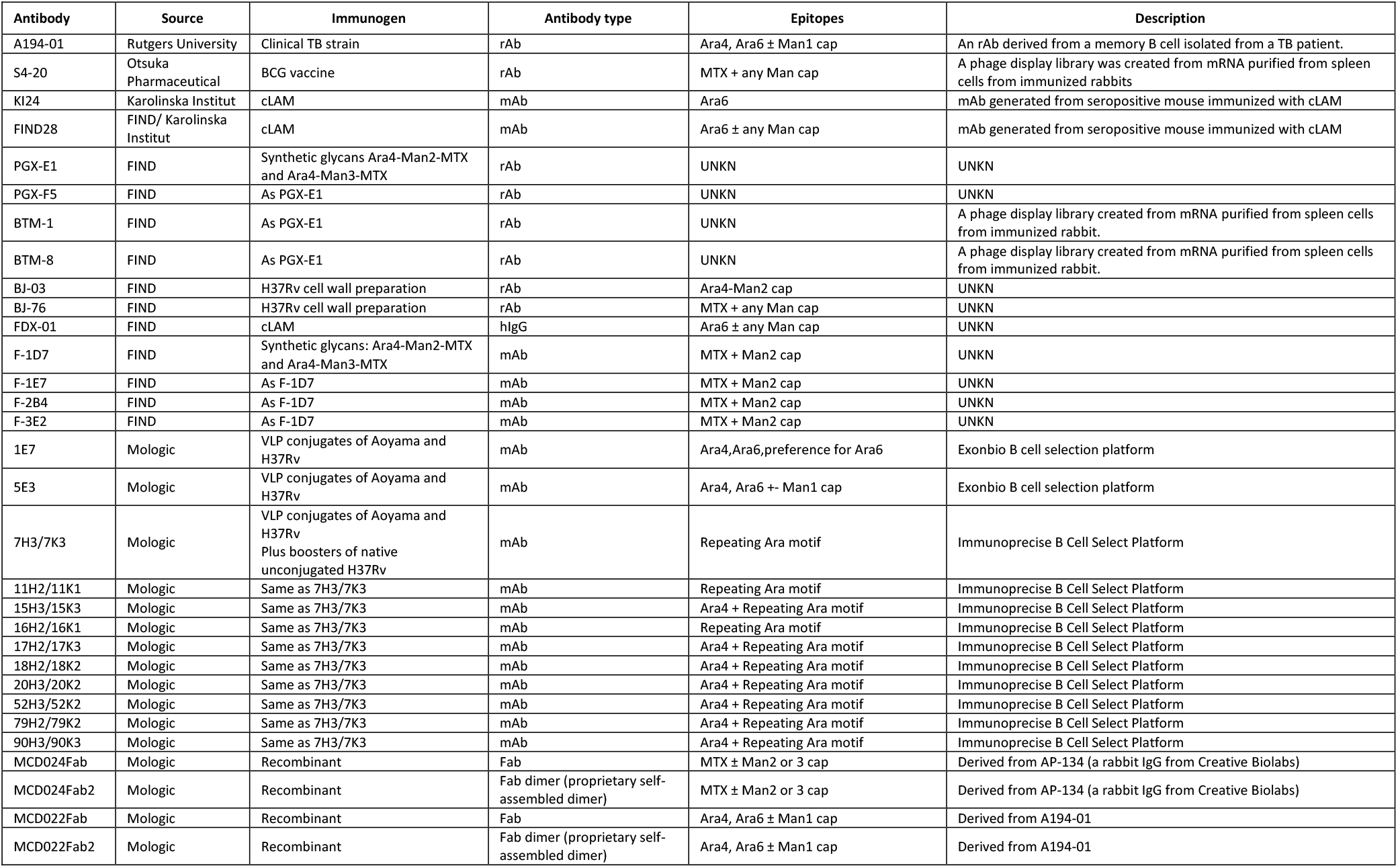
A description of the candidate Igs used in this work. Also listed is the type of antigen used for the immunogen, the type of antibody, the epitopes recognized (where known) and other pertinent information. Key: TB, tuberculosis; rAb, recombinant antibody; Ara, arabinose; Man, Mannose; MTX, 5-methylthio-xylofuranose; mIg, monoclonal antibody; UNKN, unknown; cLAM, cultured lipoarabinomannan; VLP, virus-like particle; Fab, antigen-binding fragment.

### Clinical urine samples

Clinical urine specimens were provided by FIND (Geneva, Switzerland) and Makerere University School of Medicine. All samples were collected with informed consent of the participants and using institutional review board (IRB) approved protocols. Samples from Peru, South Africa, and Vietnam were acquired from the FIND biobank. These were collected from studies with IRB approvals by the Human Research Ethics Committee of the University of Cape Town (Cape Town, South Africa), the City of Cape Town (Cape Town, South Africa; ref. 10364a); the Universidad Peruana Cayetano Heredia (Lima, Peru), and the Peruvian Ministry of Health (Lima, Peru; ref. 18829-2016); and finally the Pham Ngoc Thach Provincial Lung Hospital (Ho Chi Minh City, Vietnam), and the Vietnamese Ministry of Health (Hanoi, Vietnam; ref. 2493/QDBYT). The samples from Uganda were acquired from Global Health Labs (Bellevue, USA) and were originally collected by the Makerere University School of Medicine and the University of California, San Francisco (UCSF) with IRB approvals from the Makerere University School of Medicine Research and Ethics Committee (No. 2017-020), the Uganda National Council for Science and Technology (No. HS2210), and the UCSF Committee on Human Research (No. 17-21466). All samples were analyzed with a reference immunoassay validated in the PATH laboratory to quantify the amount of uLAM using a previously described method (33).

### An Electrochemiluminescent Immunoassay Platform for Screening Optimal Ig pairs

Each Ig was labeled to serve in the capture and detector positions, respectively. Each Ig was then assessed using a sandwich immunoassay format hosted on a highly sensitive multiplex instrument creating a total of 841 Ig pairs that were compared for their ability to detect LAM. Per their protocols, two aliquots of each antibody (1 mg/mL) were labelled with biotin (EZ-Link Sulfo-NHS-LC-Biotinylation Kit, ThermoFisher Scientific, Waltham, MA, USA) for the capture position and SULFO-TAG (GOLD SULFO-TAG NHS-Ester, MSD, Rockville, MD, USA) for detector position. Unbound biotin or SULFO-TAG was removed using Zeba spin desalting columns (ThermoFisher Scientific), and the incorporation ratio for each label was measured (See Dataverse file). Briefly, the concentration of biotinylated Igs after desalting was measured at 280 nm via spectrophotometer (Nanodrop ND-1000, ThermoFisher Scientific); biotin incorporation was measured using the Pierce Biotin quantitation kit (ThermoFisher Scientific). For measuring the incorporation of the SULFO-TAG, the protein concentration was estimated using the bicinchoninic acid (BCA) protein assay (ThermoFisher Scientific), and the SULFO-TAG label spectrophotometrically measured at 455 nm.

The biotinylated capture Igs were coupled to U-PLEX plates (MSD) via biotin-streptavidin binding to U-PLEX linkers (MSD). To prepare the capture component of the antibody array, up to 10 antibody-linker conjugates were combined in U-PLEX stop buffer at a concentration of 0.29 μg/mL per antibody, and 50 μL of this pooled mixture was added to individual well of each plate. The plates were incubated for 1 hour with shaking at 500 rpm to allow the antibody array to self-assemble to the complimentary antibody linker-binding sites on the bottom of the U-PLEX plate. Unbound material was removed by washing 3 times with 300 μL/well of phosphate buffered saline + 0.05% Tween 20 (PBS-T, pH 7.5) using a BioTek 405 TS microplate washer (BioTek Instruments Inc., Winooski, VT, USA).

Firstly, 25 μL of Buffer 22 (MSD) was added to each well in the plate before adding 25 μL of either cLAM or clinical specimens. Appropriate dilutions of cLAM were prepared in PBS + 1% BSA. The plates were incubated by shaking at 500 rpm for 1 hour at room temperature. Plates were washed 3 times with PBS-T and then 25 μL of 2 μg/mL SULFO-TAG detection antibody in Diluent 3 (MSD) was added to each well, and then incubated by shaking at 500 rpm for 1 hour at room temperature. Plates were then washed 3 times to remove excess detection reagent, and each well filled with 150 μL of 2X read buffer T (MSD). The plates were read by MESO QuickPlex SQ 120 plate reader (MSD) and the ECL from each individual array spot was subsequently measured using the Discovery Workbench v4 software (MSD).

The Ig pair S4-20/ A194-01 (capture/detector) was used as the reference pair (33) and to which each of the array spots in each well representing different capture Igs was compared. The signal-to-noise ratio (S/N) of the reference pair was expressed as 100% and each of the array spots in each plate expressed as percentile of this value. When serial dilutions of the Aoyama-B LAM were used to generate a calibration curve, the relationship of ECL to TB LAM concentration was fitted to a four-parameter logistic (4-PL) function. The limit of detection (LoD) was calculated from the fitted curve. The uLAM concentrations in clinical urine specimens were calculated by back-fitting the ECL to the 4-PL fit of a standard curve generated from serial dilutions of cLAM from the Aoyoma-B strain. All test data from this study can be publicly accessed at Dataverse (https://doi.org/10.7910/DVN/C9WVPK).

### Ig pair screening and down selection

Igs were screened in both capture and detector configurations in a matrixed format. The optimal Ig pairs for capture and detection of the TB LAM were determined over three rounds of screening. Firstly, all new Igs were screened in a matrix format using 5 ng/mL of cLAM from 5 strains of MTB complex representing 6 phylogenetic lineages (L1, L2, L3, L4, and L5:L6), and urinary LAM (uLAM) from a single TB-positive patient as target antigens. The S/N was calculated per Ig pair, and then normalized and expressed as percentage of the reference pair. The Ig pairs that recorded 90% or greater S/N than that of the reference pair on each of the five cLAMs and uLAM were selected for further screening. Other parameters used for down-selection included signal of blank/background (ideally ECL <500), and that the signal with cLAM and uLAM (ECL >10,000 at 5 ng/mL). Next, the best in class Ig pair candidates were evaluated using 7-point dilutions of the Aoyama-B LAM in 1% BSA in PBS (ranging from 40,000 to 0.016 pg/mL) in duplicate. Ig pairs were ranked in terms of their respective limit of detection (LoD). Finally, the optimal pairs were evaluated for their ability to detect uLAM in 16 clinical urine specimens collected from 4 geographical regions (Peru, South Africa, Vietnam, and Uganda).

## Results

A total of 841 Ig pairs were interrogated and compared for their ability to detect LAM. Each pair was interrogated using 5 ng/mL of *in vitro* cLAM antigen per test, which is the mid-range concentration used in the 7-point LAM calibration curve. The cLAM from MTB lineages was used to investigate if any of the candidate Ig pairs had preferential binding to discrete MTB lineages (See Table 2). The best in class candidates were pairs with a signal intensity of 90% or greater as compared to the reference pair; the pooled S/N data is shown in Table 2. The signal-minus-noise (S-N) data from lineage 1 had two capture candidates (5E3 and 1E9) that were greater than the reference pair; these were both paired with A194-01 giving values of 109% and 118%, respectively (S1 Table). Scoring these with S/N, there were 9 pairs that had a score of ≥90% (Table 2; S2 Table). The highest percentile scores were with MCD024 Fab (266%) and MCD024 Fab2 (173%) when paired with A194-01 as detector. When using S4-20 as the detector there were four candidate pairs identified, MCD024 Fab2 (90%), S4-20 (96%), MCD024 Fab (106%) and BJ-03 (109%) respectively. MCD024 Fab paired with KI24 gave a value of 120%.

**Table 2.**
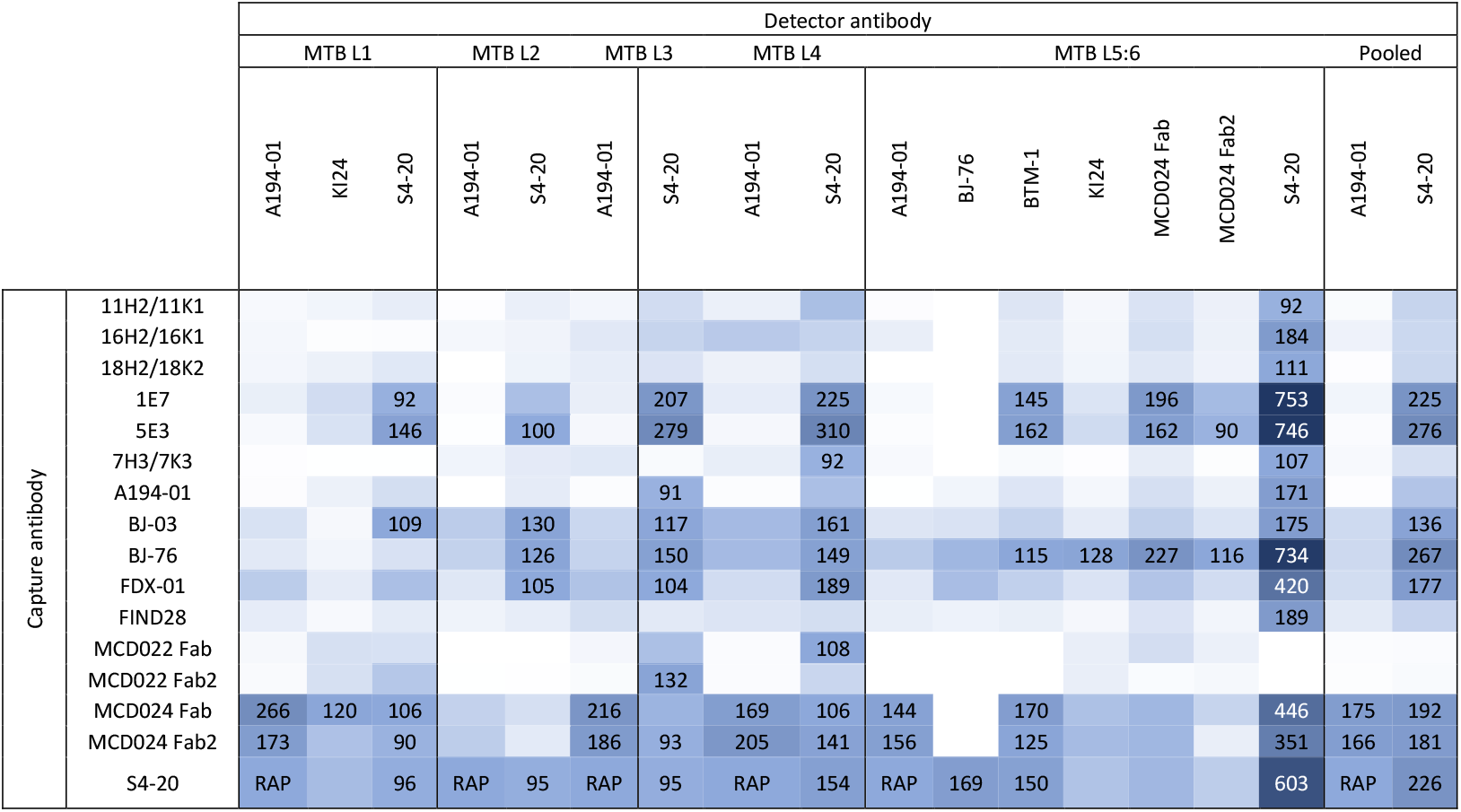
A heatmap of the normalized signal to noise (S/N) ratios used to identify the anti-LAM Ig pairs most strongly reactive to purified cLAM purified from five different lineages of MTB and the pooled data from all lineages. All data points highlighted in deepening shades of blue are ≥90% of the output from the reference assay pair (RAP). Capture Igs are in the vertical column and detector ones in the horizontal row. Unfilled boxes represent samples with a S/N output of ≤89% as compared to the RAP. Key: MTB, *Mycobacterium tuberculosis*; L, Lineage; RAP, Reference Assay Pair (S4-20/A194-01).

With lineage 2 cLAM there were no candidate Ig pairs using the S-N ratio but when S/N was applied there were 4 pairs that were ≥100% and a further one at 95%, all used S4-20 in the detector position (Table 2, S3 and S4 Tables). The capture Igs were S4-20 (95%), 5E3 (100%), FDX-01 (105%), BJ-76 (126%) and BJ-03 (130%) respectively. With MTB lineage 3 there were five pairs that had a S-N over 90% or greater (S5 Table). Here, the pair 5E3/S4-20 gave 99% and the other Igs each used A194-01 as the detector. The capture Igs were MCD022 Fab (112%), FDX-01 (121%), 1E7 (157%) and 5E3 (224%). When S/N was applied the number of candidates increased to 11 pairs but conversely none of the S-N candidates in the screen were observed when S/N was applied (Table 2, S6 Table). The increase in pairs was observed with S4-20 primarily being the detector included A194-01 (91%), MCD024 Fab2 (93%), S4-20 (95%), FDX-01 (104%), BJ-03(117%), MCD022 Fab2 (132%), BJ-76 (150%), 1E7 (207%) and 5E3 (279%). Two pairs with A194-01 as detector were >90%, MCD024 Fab2 (186%) and MCD024 Fab (216%). The S-N data for lineage 4 was very similar to lineage 1 with two only pairs, 1E7/A194-01 (104%) and 5E3/A194-01 (133%) were more reactive than the RAP (S7 Table). Using S/N there were 12 pairs that were scored (Table 2; S8 Table). Two with an A194-01 detector, MCD024 Fab and MCD024 Fab2 produced scores of 169% and 205% respectively. With S4-20 as detector there were 10 Ig pairs with MCD024 Fab and MCD024 Fab2 present in both sets. In order these were 7H3/7K3 (92%), MCD024 Fab (106%), MCD022 Fab2 (108%), MCD024 Fab2 (141%), BJ-76 (149%), S4-20 (154%), BJ-03 (161%), FDX-01 (189%), 1E7 (225%) and 5E3 (310%) respectively.

With cLAM from lineage 5:6 there were a total of 8 Ig pairs identified using S-N, one pair with A194-01 as detector, 5E3 (94%) and the remainder with S4-20 as detector in ascending order from FDX-01 (94%), MCD024 Fab2 (129%), MCD024 Fab (133%), 1E7 (174%), 5E3 (191%), S4-20 (195%) and BJ-76 (208%) (S9 Table). Using S/N, the lineage 5:6 gave the greatest number of more reactive Ig pairs with 29 pairs (excluding the RAP) overall being identified with four new Igs as the detector but all detector Igs having been previously observed with the other lineages (Table 2, S10 Table). There were BJ-76, BTM-1, MCD024 Fab and MCD024 Fab2. BJ-76 worked well in the capture position with 5 pairs being identified as opposed to just one with lineages 2-4. The use of S4-20 as the detector was also interesting in that it produced 14 pairs including the three that had the highest S/N ration in the entire set; BJ-06/S4-20 (734%), 5E3/S4-20 (746%), 1E7/S4-20 (753%) (Table 2). Overall, the data from lineages 1-4 were highly similar with A194-01 and S4-20 being the primary detectors with KI24 a sole addition for lineage 1. As noted, MTB lineage 5:6 is interesting in that a greater number of pairs were reactive, however the other Igs observed to be reactive across all of the lineages were also captured in this set. The S-N and S/N data from each lineage were pooled into one set (Table 2, S11 and S12 Tables) and the pairs with the highest S/N compared to the lineage specific pairs (Table 2). Here only A194-01 and S4-20 were consistently observed as the best detectors. With A194-01 there were only two capture Igs that were scored highly out with the reference control pair. These were MCD024 Fab (175%) and MCD024 Fab2 (166%) which are both recombinant forms derived from A194-01 recognizing Ara4, Ara6 ± Man1 cap and 1E7 and 5E3 also recognize a similar Ara4, Ara6 epitope (Table 1). With S4-20 as the detector there were eight candidate Ig pairs with a high S/N ratio, 1E7 (225%), 5E3 (276%), BJ-03 (136%), BJ-76 (267%), FDX-01 (177%), MCD024 Fab (192%), MCD024 Fab2 (181%) and S4-20 (226%). The two MCD024Fabs are both recombinant derivatives from AP-134 (Table 1).

Both A194-01 (Ara4 Ara6 Man1 cap) and S4-20 (5-methylthio-xylofuranose [MTX] ± Man cap), were the predominant detector Igs for the top performing pairs across all linages with S4-20 representing 43/74 pairs (58%) and A194-01 with 8/74 (10.8%) across the 5:6 lineages. It is also interesting to note that the MTX cap recognized by S4-20 (Table 1) is a unique epitope in LAM and so it is surprising that the self-paired S4-20 assay scored highly with each lineage. KI24 was the only other detector highlighted outside the lineage 5:6 screen 1/74 [0.14%]). As noted, the lineage 5:6 derived cLAM produced the highest number of candidate pairs yet with the S4-20 detector most prominent and BTM-1 next with 8/74 pairs (10.8%); both Igs targeting the MTX epitope. With the capture Igs there were several with similar performance including the recombinant forms MCD024Fab and MCD024Fab2, both derived from AP-134 (also targeting MTX), and these constituted 3 of the 5 top capture Igs in the screen (9/74 [12.2%], 9/74 [12.2%]and 7/74 [9.5%] respectively). BJ-76 another Ig in this group with 8/74 (10.8%) pairs having a high S/N. Each of these Igs recognize the MTX ± Man cap epitopes (see Table 1). Other Igs that had relatively better performance included 5E3 and 1E7 (7/74 [9.5%] and 6/74 [8.1%]), these are both recombinant forms that target the Ara4 and Ara6 termini. Interestingly, FDX-01 a recombinant form of FIND28 was found in 4 pairs while FIND28 was isolated only in one, both Igs targeting Ara6 ± any Man cap (Table 1).

### Screening Ig performance with uLAM

Alternative structural modifications to LAM are thought to occur during *in vivo* culture of MTB (e.g. uLAM) and an understanding of the structural differences between *in vivo* and *in vitro* derived LAM is still very limited (36,37). One study on infected mouse lung tissue suggests the succinylation of the Ara4 residue in the arabinan terminal motif and that Man caps and terminal Ara motifs may be altered or cleaved in cLAM wherein uLAM had a decrease in the terminal Ara motif and Man caps while the Ara 5 termini increased (37). In light of this proposed variance, the set of 841 Ig pairs were screened using uLAM via a single sample of urine to interrogate the performance of each Ig pair via the percentiles of the S-N and S/N ratios as compared to the RAP (S4-20/A194-01; Table 3, S13 and S14 Tables). The S/N ratios were again used to identify candidates with elevated values.

**Table 3.**
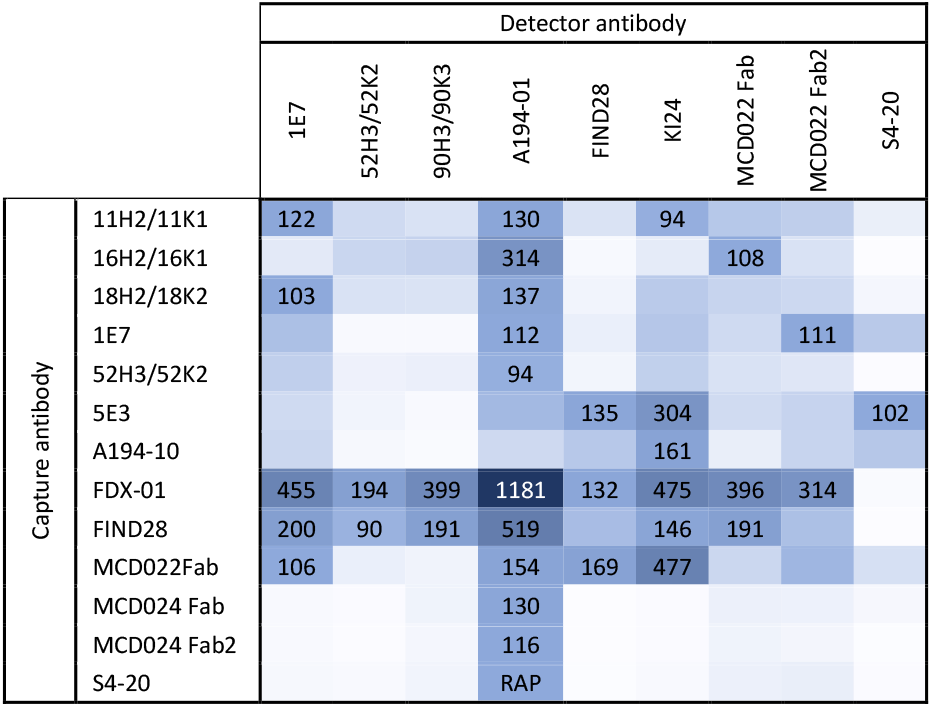
A heat map of the normalized S/N from anti-LAM Ig pairs that were most reactive to urinary LAM from a clinical sample. All highlighted data points shown are percentiles of S/N with ≥90% of the output from the reference assay Ig pair (RAP, 100%). Capture Igs are in the vertical column and detector Igs in the horizontal row. Unfilled boxes represent samples with an S/N ratio of ≤89% as compared to the RAP).

The uLAM antigen panel produced a greater number of Ig pairs with a S/N of ≥90% than the cLAM screen and represented 34 different pairs of 13 capture and 9 detector Igs in addition to the highest S/N value of 1181% (FDX-01/A194-01). Therefore the uLAM screen had some marked differences as compared to the cLAM samples (Table 3). Of the 13 Igs in the capture format, 11 were also identified in the cLAM screen (11H2/11K1, 16H2/16K1, 18H2/18K2, 1E7, 5E3, A194-01, FDX-01, FIND28, MCD024Fab, MCD024Fab2, S4-20), four in the cLAM screen were absent from uLAM (7H3/7K3, BJ-03, BJ-76, and MCD022Fab2) and two were only observed in the uLAM screen only (52H3/52K2, and MCD022Fab). For the detector format there were A194-01, KI24 and S4-20 present in both cLAM and uLAM screens. Five from the cLAM screening were absent (BJ-76, BTM-1, KI24, MCD024Fab, and MCD024Fab2) and six others were unique to uLAM (1E9, 52H3/52K2, 90H3/90K3, FIND28, MCD022Fab, and MCD022Fab2). The most significant change observed was that S4-20, the best performing detector candidate with the cLAM antigen, was now seen in only one highly reactive Ig pair when challenged with uLAM. Conversely, A194-01 was a poorer performer with cLAM but now performed as the best detector Ig for uLAM with 10/34 (29.4%) of the most reactive Ig pairs. In the capture Ig position there were 12 candidates, of which 3 represented over 50% of all the Ig pairs (FDX-01, FIND28 and MCD022 Fab [Table 3]). FDX-01 is a recombinant form of FIND28 and so it is unsurprising that both Igs exhibited similar performance in recognizing the epitopes (8/34 and 6/34 pairs respectively). However, MCD022 Fab (4/34 pairs), is an rAb derived from S4-20 and so it is unusual that the parent performed less well (Table 3).

Overall, 10 Ig pairs exhibited higher S/N ratios than the reference pair using cLAM prepared from different MTB lineages as the antigen (Table 3, Tables S1-S10), while there are 34 Ig pairs with higher S/N ratio than the reference pair when using uLAM (see Table 3, Table S12). These Ig pairs were pooled and ranked based upon their ECL signal intensities as observed when assaying both cLAM and uLAM samples (See Lineage and uLAM data folder in Dataverse file); an ECL of ≥10,000 was the threshold for strong positives, while the ECL threshold for background intensity was 500 (5X greater than the reference pair). In total, 28 Ig pairs were identified with 13 capture and 6 detector Igs with 1E7, A194-01, MCD022 Fab, and MCD022Fab2 being the only Igs that were present in both configurations (see Table 4). The pairs generating high signals with both cLAM and uLAM included the S4-20/A194-01 and FIND28/A194-01 as previously observed (33). With the panel representing both cLAM and uLAM, A194-01 and KI24 were the most common detectors with four pairs each and FDX-01 was the most common capture antibody found in 4 pairs. The capture Igs were more diverse in this panel with only MCD024 Fab and MCD024 Fab2 being represented twice.

**Table 4.** The final assessment of the optimal capture and detector Ig pairs with ranking according to their sensitivity derived from testing a clinical panel. The data includes the epitopes recognized by the Igs where known, the LAM target(s) recognized by the pairs, the limit of detection to the Aoyama B strain cLAM (MTB Lineage 2) and the performance results after screening with a panel of 16 clinical specimens and a further 8 in the case of the top performing five Ig pairs. Key: Ig, antibody; Fab, antigen-binding fragment; MTX, 5-methylthio-xylofuranose; Ara, arabinose; Man, mannose; LAM, lipoarabinomannan; C, cultured LAM; U, urinary LAM; LoD, limit of detection; Sens., senstivity; Spec., specificity.

| Capture Ig | Epitope | Detector Ig | Epitope | LAM Target | LoD (pg/mL) | Sens. (%) |  | Spec. (%) |  |
| --- | --- | --- | --- | --- | --- | --- | --- | --- | --- |
|  |  |  |  |  |  | 16 | 24 | 16 | 24 |
| S4-20 | MTX ± any Man cap | A194-01 | Ara4, Ara6 ± Man1 cap | U/C | 3.3 | 90 | 94 | 83 | 86 |
| MCD024 Fab | MTX ± Man2 or 3 cap | A194-01 | Ara4, Ara6 ± Man1 cap | C | 12.9 | 80 | 88 | 100 | 100 |
| FIND28 | Ara6 ± any Man cap | A194-01 | Ara4, Ara6 ± Man1 cap | U/C | 15.9 | 80 | 88 | 67 | 71 |
| BJ-76 | MTX ± any Man cap | A194-01 | Ara4, Ara6 ± Man1 cap | C | 5.6 | 80 | 88 | 50 | 57 |
| FDX-01 | Ara6 ± any Man cap | A194-01 | Ara4, Ara6 ± Man1 cap | U/C | 4.3 | 80 | 88 | 0 | 14 |
| 1E7 | Ara4, Ara6 | MCD022 Fab2 | Ara4, Ara6 ± Man1 cap | U/C | 10.4 | 70 | NT | 83 | NT |
| MCD022 Fab | Ara4, Ara6 ± Man1 cap | KI24 | Ara6 | U/C | 30.1 | 60 | NT | 83 | NT |
| FDX-01 | Ara6 ± any Man cap | MCD022 Fab2 | Ara4, Ara6 ± Man1 cap | U | 75.4 | 60 | NT | 83 | NT |
| 16H2/16K1 | Repeating Ara motif | A194-01 | Ara4, Ara6 ± Man1 cap | U/C | 4 | 60 | NT | 17 | NT |
| MCD024 Fab2 | MTX ± Man2 or 3 cap | A194-01 | Ara4, Ara6 ± Man1 cap | C | 6.4 | 50 | NT | 100 | NT |
| FDX-01 | Ara6 ± any Man cap | 90H3/90K3 | Ara4 | U | 171.2 | 50 | NT | 100 | NT |
| 5E3 | Ara4, Ara6 +- Man1 cap | S4-20 | MTX ± any Man cap | C | 5.4 | 40 | NT | 100 | NT |
| 1E7 | Ara4, Ara6 | S4-20 | MTX ± any Man cap | C | 3.5 | 40 | NT | 100 | NT |
| MCD02 Fab2 | Ara4, Ara6 ± Man1 cap | S4-20 | MTX ± any Man cap | C | 11.2 | 40 | NT | 100 | NT |
| FDX-01 | Ara6 ± any Man cap | 1E7 | Ara4, Ara6 | U/C | 9.6 | 40 | NT | 100 | NT |
| FDX-01 | Ara6 ± any Man cap | MCD0022 Fab | Ara4, Ara6 ± Man1 cap | U/C | 134.4 | 40 | NT | 83 | NT |
| FDX-01 | Ara6 ± any Man cap | KI24 | Ara6 | U/C | 32.1 | 30 | NT | 100 | NT |
| FIND28 | Ara6 ± any Man cap | MCD0022 Fab | Ara4, Ara6 ± Man1 cap | U | 236.5 | 30 | NT | 100 | NT |
| 5E3 | Ara4, Ara6 | KI24 | Ara6 | U/C | 18.8 | 20 | NT | 100 | NT |
| A194-01 | Ara4, Ara6 ± Man1 cap | KI24 | Ara6 | U/C | 60.2 | 20 | NT | 100 | NT |
| FDX-01 | Ara6 ± any Man cap | S4-20 | MTX ± any Man cap | C | 5.9 | 10 | NT | 100 | NT |
| MCD024 Fab | MTX ± Man cap | S4-20 | MTX ± any Man cap | C | 10 | 10 | NT | 100 | NT |
| S4-20 | MTX ± any Man cap | S4-20 | MTX ± any Man cap | C | 6.4 | 10 | NT | 100 | NT |
| BJ-76 | MTX ± any Man cap | S4-20 | MTX ± any Man cap | C | 5.8 | 10 | NT | 100 | NT |
| FIND28 | Ara6 ± any Man cap | KI24 | Ara6 | U | 256.6 | 10 | NT | 100 | NT |
| BJ-03 | Ara4-Man2 cap | A194-01 | Ara4, Ara6 ± Man1 cap | C | 4.3 | 10 | NT | 67 | NT |
| MCD024 Fab2 | MTX ± Man2 or 3 cap | S4-20 | MTX ± any Man cap | C | 23.3 | 0 | NT | 100 | NT |
| BJ-03 | Ara4-Man2 cap | S4-20 | MTX ± any Man cap | C | 28 | 0 | NT | 100 | NT |

The Ig pairs exhibited a range of LoDs spanning from 256.6 pg/mL (FIND28/KI24) to the lowest at 3.3 pg/mL (the reference pair, S4-20/A194-01). However, a further 12 Ig pairs had an LoD of ≤10 pg/mL. Therefore, while none of the new candidates in this assessment had a better analytical senstivity than the reference assay when using Aoyoma B cLAM, there are a variety of other Ig pairs with similar levels of performance. Within these Igs pairs there were no clear best in class but the epitopes recognized were roughly split with 7 capture Igs targeting Ara4 or Ara6 or Ara4, Ara6 termini (Table 4). With the detector Ig, it was close between Igs A194-01 (8 pairs) and S4-20 (9 pairs) for detector. The epitopes recognized here are either Ara4, Ara6 ± Man1 cap or a MTX ± Man cap, respectively.

The performance of the 28 Ig pairs were compared on a panel of 16 clinical samples from confirmed TB positive and negative cases and for participants who were also diagnosed as HIV positive or negative. The diagnostic sensitivity and specificity were calculated for each pair from the clinical data (molecular and culture diagnosis). When using sensitivity as the key metric, the S4-20/A194-01 pair was again best in class at 90%. Four other pairs had a sensitivity of 80% with each using A194-01 as the detector (Table 4). The associated capture antibodies were MCD024 Fab, FIND28, BJ-076 and FDX-01. When a further 8 clinical samples were screened with these 5 pairs the sensitivity improved for each of them but with the order of ranking not changing. However, when specificity was also assessed the performance of the top 5 pairs became varied. The ranking changed slightly with MCD024 Fab/A194-01 having 100% specificity while S4-20/A194-01, was next at 84%. The MTX motif recognized by S4-20, MCD024 Fab and BJ-76 is unique to MTB and yet BJ-76 was more indiscriminate in its binding with a specificity of 56% as opposed to S4-20 and MCD024 Fab.

## Discussion

A key step in the cascade of TB care is facilitating better and earlier access to rapid screening for infection. The relative simplicity of collecting a urine sample for the detection of uLAM antigens makes rapid antigen assays a candidate to fulfil this role. However, to be of universal value, the sensitivity of the uLAM assay must be improved to meet the ranges reflected in the target product profile prepared by the WHO in both HIV positive and negative populations (9). Several immunoassays to detect LAM in either urine and sputum have been developed (23,24,32,33,35,38) including the Alere LF-LAM Determine assay (Abbott Diagnostics), currently the only commercially available test for uLAM with WHO policy guidelines (39,40). These guidelines reflect the clinical sensitivity challenge when using the Determine assay to diagnose active TB as the assay has insufficient sensitivity to detect active TB in many HIV negative cases (38) or with TB cases having CD4 count >100 cells/mL. There are also challenges based on which LAM epitopes are displayed when MTB cells are cultured *in vivo* (i.e. by infection) or *in vitro* (in growth media) and also structural differences within different TB lineages (36,37). Other considerations include what is the absolute concentration range of uLAM in active TB cases, and if confounders can mask LAM in urine (e.g., proteinuria, urinary tract infections) (41,42). The limited performance of the Determine assay may be attributed to the performance of the Ig pair used in addition to the small sample volume (60 μL) that limits the sensitive detection of uLAM (e.g. <250 pg/mL). We aimed to identify candidate Ig pairs from new antibody development efforts to improve the sensitivity of rapid antigen assays targeting uLAM. We accessed a range of Igs, in addition to their derivatives as Fabs from the light and heavy chain variable regions. The immunogens used were whole MTB cells from *in vitro* culture, purified cLAM, and synthetic glycans aimed at specific epitopes, e.g. the MTX motif (Table 1).

A highly versatile microarray platform enabled the rapid screening of 841 candidate pairs in both the detection and candidate orientations using a variety of epitopes to equivocally assess performance. Overall, the results from these rounds of screening did not identify any new candidate Igs with better performance. Somewhat surprisingly given the number of LAM Igs pairings assessed, the reference pair S4-20/A194-01, remained the best overall candidate. Recombinant derivatives of A194-01 were assessed and while reaching the top tier after three rounds of screening they did not perform as well as the original candidates with uLAM. This may be attributed to subtle structural changes when expressed with a different core antibody structure or lost performance with events such as misfolding during production that can significantly affect performance (43).

An interesting outcome of this work was the demonstration that some Ig pairs each had good performance when using cLAM antigen derived from *in vitro* cultured TB cells but their performance on uLAM samples was poorer. This suggests that uLAM may have structural differences from cLAM which is a homogenous and relatively pure material. However, uLAM is generated from *in vivo* culture in the host and there is preliminary evidence of structural differences with succinylation and a higher prevalence of Ara 5 and a profound lack of Ara6 in the urine of a HIV-ve/TB +ve subject (19,37). We speculate that there are opportunities for modification of uLAM termini via enzymatic or physiochemical decay removing the MTX-1, Man caps or Ara termini as shown with an increase of Ara5 in uLAM (37). The heterogeneity of uLAM means that the detailed composition and structure of uLAM remains unresolved as compared to cLAM. Other compounds found in urine may also affect Ig binding via the masking of epitopes by proteins or other materials sequestering them or other compounds binding to the Igs, rendering them non-or partially functional. Despite the considerable amount of effort in generating Igs to LAM to date, until their screening is focused on performance assessments using uLAM sources, Igs resulting in improved performance are unlikely to be found.

We used cLAM from MTB strains representing 6 different lineages in our screening process and with four lineages observed little difference and with the 5-6 lineage many more Ig pairs were reactive. This suggests that some epitopes may be more accessible in this lineage to but overall the majority of these Ig pairs then performed poorly with uLAM. In looking at development of the Igs used in this study, the majority (19) used cLAM or whole TB cells as the immunogen. As such, if novel structural LAM epitopes are created only by *in vivo* cultured cells then these potential epitopes will be missed and conversely, if recombinant Igs are screened from the memory B cell population of convalescent TB patients (e.g. A194-01, (22)), using cLAM as the target antigen in screening may miss Igs targeting uLAM epitopes not present in cLAM. Ig development efforts using synthetic glycans to generate Igs to the TB specific MTX-1 motif, targeted by S4-20 (23) did not produce a successful product but highlight the potential for targeted Ig generation to epitopes. Using glycan arrays has played a key role in understanding which LAM epitopes are targeted by the antibody candidates to date (22).

Our study had a range of limitations. Firstly, screening with cLAM may not find the most suitable Igs that target uLAM. Our initial hypothesis was that screening with multiple TB lineages may identify some key structural differences in LAM that could be reflected by different Ig pairs having improved performance on specific lineages (36). As noted we did not see appreciable differences between lineages and using cLAM from one lineage for further down select only selected for those Igs that had better performance with cLAM. As such, a larger sample set of antibody pairs should have been screened on more urine samples to identify if the initial screens using cLAM was inadequate. Our uLAM samples, while reflecting geographic diversity, were very small in terms of their number. A further study will focus on fewer Igs using significantly greater numbers of urine samples. A further perspective is the nature of the assay used for screening. A liquid immunoassay works optimally with Igs that have slower on rate kinetics as the Igs have sufficient time to recognize and bind to their target epitopes. In a rapid antigen test, the exposure time of the antibody to its target is likely a few seconds at most and so the kinetics of what binds best are different from a liquid immunoassay. A recent study compared a subset of Ig pairs in this work (289 pairs) via detection of Aoyoma-B cLAM on the MSD platform versus an LFA screening platform (30). In comparing signal outputs from both methods, they showed a weak positive correlation, with *R* values of 0.30 (*p* < 0.0001) and 0.36 (*p* < 0.0001) for S–N and S/N. New Igs continue to be produced, some taking novel approaches using camelid sources to develop Igs that can recognize smaller antigen targets. FIND hosts a biorepository with multiple urine samples for developers to access and PATH recently developed a method to purify and enrich up to 50% of the total uLAM from urine for early screening of antibody development candidates (44). These efforts in conjunction with ongoing innovations in sample enrichment, assay design, and signal amplification may result in the affordable and effective rapid antigen test that is so urgently needed to identify TB cases and offer them entry into the cascade of care for a curable disease.

## Acknowledgements

We are most grateful to the consented participants who gave the urine samples from the study cohorts in Ghana, Peru, South Africa, and Vietnam. We greatly appreciate the work from the staff at the Makerere University (Uganda) in collecting and processing the urine specimens and the support of Dr. Adithya Cattamanchi (UCSF) to access these samples through Drs. John Connelly and Benjamin Grant (Global Health Labs). Similarly, we thank the project leadership and study staff from the University of Cape Town (South Africa), Universidad Peruana Cayetano Heredia (Peru) and the Pham Ngoc Thach Provincial Lung Hospital (Vietnam) for their efforts in the collection of urine specimens and thank Dr. Morten Ruhwald (FIND) for providing access to these collections. We thank Ms. Jennifer Chin for her diligence in managing the Material Transfer Agreements that were necessary to access the many different sources of specimens and reagents used in this study. This publication is based on research funded by the Bill & Melinda Gates Foundation via a grant (INV-008079) to DSB. The findings and conclusions contained within are those of the authors and do not necessarily reflect positions or policies of the Bill & Melinda Gates Foundation. The research reported in this publication was also supported by the National Institute of Allergy and Infectious Diseases of the National Institutes of Health under award number RO1AI132680 to DC. DC spent a cumulative 10% of her effort on the research in characterization of LAM isolated from the lineages described, evaluating data and editing manuscript. The content is solely the responsibility of the authors and does not necessarily represent the official views of the National Institutes of Health.

## Notes

### Competing Interest Statement

James Schouten and Paul Davis are employees of Mologic Ltd., Bedford, UK. Masanori Kawasaki is an employee of Otsuka Pharmaceutical Co., Ltd., Tokyo, Japan. All other authors have no competing financial interests.

