## Supplemental tables for "Performance of novel antibodies for lipoarabinomannan to develop diagnostic tests for *Mycobacterium tuberculosis*"

### S1 Table. Lineage 1 S - N ratio vs RAP

| | | Detector antibody | | | | | | | | | | | | | | | | | | | | | | | | | | | | |
|---|---|---|---|---|---|---|---|---|---|---|---|---|---|---|---|---|---|---|---|---|---|---|---|---|---|---|---|---|---|---|
| | | 11H2/ 11K1 | 15H3/ 15K3 | 16H2/ 16K1 | 17H2/ 17K3 | 18H2/ 18K2 | 1E7 | 20H3/ 20K2 | 52H3/ 52K2 | 5E3 | 79H2/ 79K2 | 7H3/ 7K3 | 90H3/ 90K3 | A194 -01 | BJ -03 | BJ -76 | BTM -1 | BTM -8 | F-1 D7 | F-1 E7 | F -2B4 | F-3 E2 | FDX -01 | FIND 28 | KI24 | MCD 022 Fab | MCD 022 Fab2 | MCD 024 Fab | MCD 024 Fab2 | S4- 20 |
| Capture antibody | 11H2/11K1 | 0 | 0 | 0 | 0 | 0 | 2 | 0 | 1 | 4 | 0 | 0 | 2 | 18 | 1 | 1 | 2 | 1 | 0 | 0 | 0 | 0 | 0 | 1 | 8 | 5 | 4 | 3 | 1 | 6 |
| | 15H3/15K3 | 0 | 0 | 0 | 0 | 0 | 0 | 0 | 0 | 0 | 0 | 0 | 0 | 1 | 0 | 0 | 0 | 0 | 0 | 0 | 0 | 0 | 0 | 0 | 0 | 0 | 0 | 0 | 0 | 0 |
| | 16H2/16K1 | 0 | 0 | 0 | 0 | 0 | 0 | 0 | 0 | 0 | 0 | 0 | 0 | 4 | 0 | 0 | 1 | 0 | 0 | 0 | 0 | 0 | 0 | 0 | 1 | 2 | 1 | 1 | 0 | 1 |
| | 17H2/17K3 | 0 | 0 | 0 | 0 | 0 | 0 | 0 | 0 | 0 | 0 | 0 | 0 | 1 | 0 | 0 | 0 | 0 | 0 | 0 | 0 | 0 | 0 | 0 | 1 | 1 | 0 | 0 | 0 | 0 |
| | 18H2/18K2 | 0 | 0 | 0 | 0 | 0 | 3 | 0 | 1 | 4 | 0 | 0 | 1 | 24 | 1 | 1 | 2 | 1 | 0 | 0 | 0 | 0 | 1 | 1 | 7 | 5 | 3 | 3 | 1 | 8 |
| | 1E7 | 3 | 2 | 0 | 0 | 0 | 8 | 0 | 2 | 13 | 0 | 0 | 5 | 109 | 3 | 6 | 9 | 3 | 0 | 0 | 0 | 0 | 3 | 2 | 24 | 22 | 18 | 10 | 5 | 27 |
| | 20H3/20K2 | 0 | 0 | 0 | 0 | 0 | 0 | 0 | 0 | 1 | 0 | 0 | 0 | 6 | 0 | 0 | 1 | 0 | 0 | 0 | 0 | 0 | - | 0 | 2 | 1 | 1 | 1 | 0 | 2 |
| | 52H3/52K2 | 0 | 0 | 0 | 0 | 0 | 1 | 0 | 0 | 0 | 0 | 0 | 0 | 6 | 0 | 0 | 0 | 0 | 0 | 0 | 0 | 0 | 0 | 0 | 1 | 1 | 0 | 0 | 0 | 1 |
| | 5E3 | 5 | 3 | 0 | 0 | 1 | 10 | 0 | 2 | 24 | 0 | 0 | 4 | 118 | 7 | 9 | 11 | 4 | 0 | 0 | 0 | 0 | 6 | 3 | 37 | 26 | 20 | 12 | 6 | 31 |
| | 79H2/79K2 | 0 | 0 | 0 | 0 | 0 | 0 | 0 | 0 | 0 | 0 | 0 | 0 | 4 | 0 | 0 | 0 | 0 | 0 | 0 | 0 | 0 | 0 | 0 | 1 | 1 | 0 | 0 | 0 | 0 |
| | 7H3/7K3 | 0 | 0 | 0 | 0 | 0 | 0 | 0 | 0 | 0 | 0 | 0 | 0 | 1 | 0 | 0 | 0 | 0 | 0 | 0 | 0 | 0 | 0 | 0 | 0 | 1 | 0 | 0 | 0 | 0 |
| | 90H3/90K3 | 0 | 0 | 0 | 0 | 0 | 1 | 0 | 0 | 1 | 0 | 0 | 1 | 11 | 0 | 0 | 0 | 0 | 0 | 0 | 0 | 0 | 0 | 0 | 2 | 2 | 1 | 1 | 0 | 2 |
| | A194-01 | 2 | 1 | 0 | 0 | 0 | 5 | 0 | 2 | 5 | 0 | 0 | 4 | 72 | 0 | 1 | 4 | 1 | 0 | 0 | 0 | 0 | 1 | 2 | 18 | 10 | 10 | 4 | 2 | 11 |
| | BJ-03 | 2 | 1 | 0 | 0 | 0 | 5 | 0 | 0 | 12 | 0 | 0 | 1 | 46 | 2 | 4 | 6 | 2 | 0 | 0 | 0 | 0 | 1 | 1 | 3 | 8 | 6 | 5 | 2 | 32 |
| | BJ-76 | 1 | 1 | 0 | 0 | 0 | 5 | 0 | 0 | 9 | 0 | 0 | 1 | 36 | 1 | 1 | 3 | 1 | 0 | 0 | 0 | 0 | 2 | 1 | 4 | 7 | 5 | 3 | 1 | 11 |
| | BTM-1 | 0 | 0 | 0 | 0 | 0 | 0 | 0 | 0 | 1 | 0 | 0 | 0 | 4 | 0 | 0 | 0 | 0 | 0 | 0 | 0 | 0 | 0 | 0 | 1 | 1 | 1 | 1 | 0 | 2 |
| | BTM-8 | 0 | 0 | 0 | 0 | 0 | 0 | 0 | 0 | 0 | 0 | 0 | 0 | 2 | 0 | 0 | 0 | 0 | 0 | 0 | 0 | 0 | 0 | 0 | 1 | 1 | 0 | 0 | 0 | 1 |
| | F-1D7 | 0 | 0 | 0 | 0 | 0 | 0 | 0 | 0 | 0 | 0 | 0 | 0 | 0 | 0 | 0 | 0 | 0 | 0 | 0 | 0 | 0 | 0 | 0 | 0 | 0 | 0 | 0 | 0 | 0 |
| | F-1E7 | 0 | 0 | 0 | 0 | 0 | 0 | 0 | 0 | 0 | 0 | 0 | 0 | 0 | 0 | 0 | 0 | 0 | 0 | 0 | 0 | 0 | 0 | 0 | 0 | 0 | 0 | 0 | 0 | 0 |
| | F-2B4 | 0 | 0 | 0 | 0 | 0 | 0 | 0 | 0 | 0 | 0 | 0 | 0 | 0 | 0 | 0 | 0 | 0 | 0 | 0 | -4 | 0 | 0 | 0 | 0 | 0 | 0 | 0 | 0 | 0 |
| | F-3E2 | 0 | 0 | 0 | 0 | 0 | 0 | 0 | 0 | 0 | 0 | 0 | 0 | 0 | 0 | 0 | 0 | 0 | 0 | 0 | 0 | 0 | 0 | 0 | 0 | 0 | 0 | 0 | 0 | 0 |
| | FDX-01 | 1 | 1 | 0 | 0 | 0 | 6 | 0 | 1 | 8 | 0 | 0 | 4 | 77 | 1 | 2 | 3 | 1 | 0 | 0 | 0 | 0 | 1 | 2 | 8 | 11 | 6 | 4 | 1 | 14 |
| | FIND28 | 0 | 0 | 0 | 0 | 0 | 2 | 0 | 1 | 2 | 0 | 0 | 1 | 25 | 0 | 1 | 1 | 0 | 0 | 0 | 0 | 0 | 0 | 1 | 3 | 5 | 2 | 2 | 0 | 5 |
| | MCD022 Fab | 2 | 1 | 0 | 0 | 0 | 10 | 0 | 3 | 8 | 0 | 0 | 6 | 78 | 0 | 1 | 2 | 1 | 0 | 0 | 0 | 0 | 1 | 2 | 25 | 14 | 14 | 6 | 2 | 8 |
| | MCD022 Fab2 | 2 | 1 | 0 | 0 | 0 | 8 | 0 | 1 | 8 | 0 | 0 | 3 | 69 | 1 | 2 | 3 | 1 | 0 | 0 | 0 | 0 | 1 | 1 | 17 | 11 | 12 | 7 | 3 | 16 |
| | MCD024 Fab | 3 | 2 | 0 | 0 | 0 | 12 | 0 | 1 | 17 | 0 | 0 | 2 | 79 | 2 | 2 | 7 | 3 | 0 | 0 | 0 | 0 | 2 | 6 | 30 | 15 | 13 | 7 | 4 | 26 |
| | MCD024 Fab2 | 3 | 1 | 0 | 0 | 0 | 10 | 0 | 1 | 14 | 0 | 0 | 2 | 70 | 2 | 2 | 6 | 2 | 0 | 0 | 0 | 0 | 2 | 4 | 26 | 13 | 13 | 6 | 3 | 23 |
| | S4-20 | 3 | 1 | 0 | 0 | 0 | 7 | 0 | 1 | 11 | 0 | 0 | 2 | 100 | 4 | 3 | 6 | 2 | 0 | 0 | 0 | 0 | 4 | 4 | 24 | 13 | 11 | 6 | 3 | 30 |

Table S2. Lineage 1 S / N ratio vs RAP

| | Detector antibody | | | | | | | | | | | | | | | | | | | | | | | | | | | | |
|---|---|---|---|---|---|---|---|---|---|---|---|---|---|---|---|---|---|---|---|---|---|---|---|---|---|---|---|---|---|
| Capture antibody | 11H2/ 11K1 | 15H3/ 15K3 | 16H2/ 16K1 | 17H2/ 17K3 | 18H2/ 18K2 | 1E7 | 20H3/ 20K2 | 52H3/ 52K2 | 5E3 | 79H2/ 79K2 | 7H3/ 7K3 | 90H3/ 90K3 | A194 -01 | BJ -03 | BJ -76 | BTM -1 | BTM -8 | F-1 D7 | F-1 E7 | F-2 B4 | F-3 E2 | FDX -01 | FIND 28 | KI24 | MCD 022 Fab | MCD 022 Fab2 | MCD 024 Fab | MCD 024 Fab2 | S4- 20 |
| 11H2/11K1 | 1 | 0 | 0 | 0 | 0 | 6 | 0 | 2 | 7 | 0 | 0 | 2 | 8 | 2 | 3 | 6 | 3 | 0 | 0 | 0 | 0 | 2 | 3 | 13 | 7 | 7 | 13 | 5 | 22 |
| 15H3/15K3 | 0 | 0 | 0 | 0 | 0 | 0 | 0 | 0 | 0 | 0 | 0 | 0 | 1 | 0 | 0 | 0 | 0 | 0 | 0 | 0 | 0 | 0 | 0 | 1 | 1 | 0 | 0 | 0 | 1 |
| 16H2/16K1 | 0 | 0 | 0 | 0 | 0 | 1 | 0 | 1 | 2 | 0 | 0 | 1 | 10 | 0 | 1 | 2 | 1 | 0 | 0 | 0 | 0 | 0 | 0 | 3 | 5 | 5 | 3 | 1 | 4 |
| 17H2/17K3 | 0 | 0 | 0 | 0 | 0 | 1 | 0 | 0 | 1 | 0 | 0 | 0 | 1 | 0 | 0 | 0 | 0 | 0 | 0 | 0 | 0 | 0 | 0 | 1 | 2 | 1 | 1 | 0 | 1 |
| 18H2/18K2 | 1 | 0 | 0 | 0 | 0 | 9 | 0 | 1 | 9 | 0 | 0 | 2 | 11 | 2 | 3 | 5 | 2 | 0 | 0 | 0 | 0 | 3 | 4 | 17 | 7 | 7 | 8 | 5 | 28 |
| 1E7 | 4 | 2 | 0 | 0 | 1 | 14 | 1 | 1 | 10 | 0 | 0 | 1 | 20 | 10 | 20 | 25 | 11 | 0 | 0 | 0 | 0 | 13 | 8 | 41 | 13 | 21 | 45 | 20 | 92 |
| 20H3/20K2 | 0 | 0 | 0 | 0 | 0 | 2 | 0 | 0 | 2 | 0 | 0 | 1 | 10 | 0 | 0 | 2 | 1 | 0 | 0 | 0 | 0 | - | 0 | 5 | 3 | 3 | 3 | 1 | 6 |
| 52H3/52K2 | 0 | 0 | 0 | 0 | 0 | 2 | 0 | 0 | 1 | 0 | 0 | 1 | 3 | 0 | 1 | 1 | 0 | 0 | 0 | 0 | 0 | 0 | 1 | 3 | 2 | 1 | 1 | 0 | 3 |
| 5E3 | 3 | 2 | 0 | 1 | 1 | 8 | 1 | 1 | 5 | 0 | 0 | 1 | 7 | 20 | 31 | 34 | 13 | 0 | 0 | 0 | 0 | 29 | 10 | 35 | 8 | 7 | 48 | 24 | 146 |
| 79H2/79K2 | 0 | 0 | 0 | 0 | 0 | 2 | 0 | 0 | 0 | 0 | 0 | 1 | 2 | 0 | 0 | 1 | 0 | 0 | 0 | 0 | 0 | 0 | 0 | 2 | 1 | 0 | 1 | 0 | 2 |
| 7H3/7K3 | 0 | 0 | 0 | 0 | 0 | 0 | 0 | 0 | 1 | 0 | 0 | 0 | 3 | 0 | 0 | 0 | 0 | 0 | 0 | 0 | 0 | 0 | 0 | 1 | 2 | 1 | 1 | 0 | 1 |
| 90H3/90K3 | 1 | 0 | 0 | 0 | 0 | 3 | 0 | 0 | 1 | 0 | 0 | 1 | 4 | 0 | 0 | 1 | 1 | 0 | 0 | 0 | 0 | 0 | 1 | 6 | 2 | 2 | 2 | 1 | 7 |
| A194-01 | 2 | 1 | 0 | 0 | 0 | 4 | 0 | 1 | 2 | 0 | 0 | 1 | 4 | 2 | 3 | 13 | 5 | 0 | 0 | 0 | 0 | 4 | 6 | 17 | 3 | 5 | 16 | 9 | 39 |
| BJ-03 | 8 | 5 | 0 | 0 | 1 | 18 | 0 | 2 | 44 | 0 | 0 | 3 | 35 | 5 | 12 | 19 | 6 | 0 | 0 | 0 | 0 | 6 | 3 | 9 | 18 | 17 | 20 | 11 | 109 |
| BJ-76 | 4 | 5 | 0 | 0 | 1 | 21 | 0 | 1 | 25 | 0 | 0 | 2 | 26 | 4 | 4 | 9 | 3 | 0 | 0 | 0 | 0 | 7 | 3 | 12 | 20 | 13 | 14 | 4 | 35 |
| BTM-1 | 1 | 0 | 0 | 0 | 0 | 2 | 0 | 0 | 3 | 0 | 0 | 1 | 9 | 1 | 1 | 1 | 1 | 0 | 0 | 0 | 0 | 1 | 1 | 6 | 3 | 3 | 2 | 1 | 6 |
| BTM-8 | 0 | 0 | 0 | 0 | 0 | 1 | 0 | 0 | 2 | 0 | 0 | 0 | 7 | 1 | 1 | 1 | 0 | 0 | 0 | 0 | 0 | 1 | 0 | 3 | 2 | 2 | 1 | 1 | 4 |
| F-1D7 | 0 | 0 | 0 | 0 | 0 | 0 | 0 | 0 | 0 | 0 | 0 | 0 | 0 | 0 | 0 | 0 | 0 | 0 | 0 | 0 | 0 | 0 | 0 | 0 | 0 | 0 | 0 | 0 | 0 |
| F-1E7 | 0 | 0 | 0 | 0 | 0 | 0 | 0 | 0 | 0 | 0 | 0 | 0 | 0 | 0 | 0 | 0 | 0 | 0 | 0 | 0 | 0 | 0 | 0 | 0 | 0 | 0 | 0 | 0 | 0 |
| F-2B4 | 0 | 0 | 0 | 0 | 0 | 0 | 0 | 0 | 0 | 0 | 0 | 0 | 0 | 0 | 0 | 0 | 0 | 0 | 0 | 0 | 0 | 0 | 0 | 0 | 1 | 0 | 0 | 0 | 0 |
| F-3E2 | 0 | 0 | 0 | 0 | 0 | 0 | 0 | 0 | 0 | 0 | 0 | 0 | 0 | 0 | 0 | 0 | 0 | 0 | 0 | 0 | 0 | 0 | 0 | 0 | 0 | 0 | 0 | 0 | 0 |
| FDX-01 | 4 | 4 | 0 | 0 | 1 | 28 | 0 | 5 | 29 | 1 | 0 | 12 | 61 | 4 | 10 | 11 | 4 | 0 | 0 | 0 | 0 | 5 | 7 | 24 | 33 | 24 | 20 | 8 | 75 |
| FIND28 | 3 | 3 | 0 | 0 | 0 | 9 | 0 | 4 | 14 | 1 | 0 | 5 | 22 | 1 | 4 | 5 | 2 | 0 | 0 | 0 | 0 | 1 | 1 | 8 | 14 | 7 | 7 | 2 | 24 |
| MCD022 Fab | 2 | 1 | 0 | 0 | 0 | 7 | 0 | 1 | 3 | 0 | 0 | 1 | 9 | 1 | 3 | 8 | 2 | 0 | 0 | 0 | 0 | 3 | 7 | 37 | 4 | 6 | 21 | 9 | 36 |
| MCD022 Fab2 | 2 | 1 | 0 | 0 | 0 | 6 | 0 | 1 | 3 | 0 | 0 | 0 | 6 | 2 | 5 | 7 | 4 | 0 | 0 | 0 | 0 | 5 | 1 | 37 | 2 | 1 | 1 | 3 | 66 |
| MCD024 Fab | 16 | 10 | 0 | 1 | 2 | 56 | 1 | 5 | 69 | 1 | 0 | 10 | 266 | 9 | 8 | 36 | 17 | 0 | 0 | 0 | 0 | 10 | 30 | 120 | 53 | 57 | 25 | 15 | 106 |
| MCD024 Fab2 | 10 | 5 | 0 | 1 | 1 | 43 | 1 | 3 | 55 | 0 | 0 | 6 | 173 | 6 | 5 | 19 | 7 | 0 | 0 | 0 | 0 | 6 | 14 | 72 | 40 | 28 | 16 | 4 | 90 |
| S4-20 | 9 | 5 | 0 | 1 | 1 | 29 | 1 | 3 | 41 | 0 | 0 | 8 | 100 | 11 | 9 | 20 | 8 | 0 | 0 | 0 | 0 | 20 | 21 | 79 | 33 | 37 | 19 | 12 | 96 |

Table S3 Lineage 2 S - N ratio vs RAP

| | | Detector antibody | | | | | | | | | | | | | | | | | | | | | | | | | | | |
|---|---|---|---|---|---|---|---|---|---|---|---|---|---|---|---|---|---|---|---|---|---|---|---|---|---|---|---|---|---|
| | | 11H2/11K1 | 15H3/15K3 | 16H2/16K1 | 17H2/17K3 | 18H2/18K2 | 1E7 | 20H3/20K2 | 52H3/52K2 | 5E3 | 79H2/79K2 | 7H3/7K3 | 90H3/90K3 | A194-01 | BJ-03 | BJ-76 | BTM-1 | BTM-8 | F-1D7 | F-1E7 | F-2B4 | F-3E2 | FDX-01 | FIND28 | KI24 | MCD022 Fab | MCD022 Fab2 | MCD024 Fab | MCD024 Fab2 | S4-20 |
| Capture antibody | 11H2/11K1 | 1 | 0 | 0 | 0 | 0 | 2 | 0 | 0 | 3 | 0 | 0 | 1 | 13 | 0 | 17 | 3 | 1 | 0 | 1 | 0 | 0 | 10 | 1 | 3 | 2 | 2 | 2 | 1 | 9 |
| | 15H3/15K3 | 0 | 0 | 0 | 0 | 0 | 0 | 0 | 0 | 0 | 0 | 0 | 0 | 0 | 0 | 1 | 0 | 0 | 0 | 0 | 0 | 0 | 0 | 0 | 0 | 0 | 0 | 0 | 0 | 0 |
| | 16H2/16K1 | 1 | 0 | 0 | 0 | 0 | 1 | 0 | 0 | 3 | 0 | 0 | 1 | 16 | 0 | 11 | 4 | 1 | 0 | 0 | 0 | 0 | 6 | 1 | 3 | 3 | 3 | 3 | 1 | 9 |
| | 17H2/17K3 | 0 | 0 | 0 | 0 | 0 | 0 | 0 | 0 | 0 | 0 | 0 | 0 | 1 | 0 | 0 | 0 | 0 | 0 | 0 | 0 | 0 | 0 | 0 | 0 | 0 | 0 | 0 | 0 | 1 |
| | 18H2/18K2 | 0 | 0 | 0 | 0 | 0 | 1 | 0 | 0 | 2 | 0 | 0 | 0 | 11 | 0 | 1 | 2 | 1 | 0 | 0 | 0 | 0 | 0 | 0 | 2 | 2 | 2 | 2 | 2 | 7 |
| | 1E7 | 5 | 3 | 0 | 0 | 0 | 8 | 0 | 1 | 10 | 0 | 0 | 2 | 64 | 1 | 63 | 15 | 5 | 1 | 3 | 0 | 1 | 33 | 2 | 9 | 7 | 8 | 8 | 4 | 43 |
| | 20H3/20K2 | 0 | 0 | 0 | 0 | 0 | 0 | 0 | 0 | 1 | 0 | 0 | 0 | 2 | 0 | 0 | 1 | 0 | 0 | 0 | 0 | 0 | - | 0 | 1 | 1 | 0 | 1 | 1 | 2 |
| | 52H3/52K2 | 0 | 0 | 0 | 0 | 0 | 0 | 0 | 0 | 0 | 0 | 0 | 0 | 2 | 0 | 0 | 0 | 0 | 0 | 0 | 0 | 0 | 0 | 0 | 0 | 0 | 0 | 0 | 1 | 1 |
| | 5E3 | 8 | 4 | 0 | 0 | 1 | 12 | 0 | 2 | 15 | 0 | 0 | 3 | 80 | 1 | 84 | 20 | 8 | 1 | 4 | 1 | 1 | 42 | 4 | 14 | 9 | 10 | 10 | 6 | 52 |
| | 79H2/79K2 | 0 | 0 | 0 | 0 | 0 | 0 | 0 | 0 | 0 | 0 | 0 | 0 | 1 | 0 | 0 | 0 | 0 | 0 | 0 | 0 | 0 | 0 | 0 | 0 | 0 | 0 | 0 | 2 | 0 |
| | 7H3/7K3 | 0 | 0 | 0 | 0 | 0 | 1 | 0 | 0 | 1 | 0 | 0 | 0 | 19 | 0 | 5 | 1 | 0 | 0 | 0 | -1 | 0 | 3 | 1 | 1 | 1 | 1 | 1 | 0 | 12 |
| | 90H3/90K3 | 0 | 0 | 0 | 0 | 0 | 0 | 0 | 0 | 0 | 0 | 0 | 0 | 3 | 0 | 0 | 0 | 0 | 0 | 0 | 0 | 0 | 0 | 0 | 0 | 1 | 0 | 0 | 4 | 1 |
| | A194-01 | 2 | 1 | 0 | 0 | 0 | 3 | 0 | 0 | 3 | 0 | 0 | 1 | 22 | 1 | 2 | 4 | 1 | 0 | 0 | 0 | 0 | 1 | 1 | 4 | 2 | 3 | 3 | 1 | 12 |
| | BJ-03 | 6 | 4 | 0 | 0 | 1 | 10 | 0 | 1 | 17 | 0 | 0 | 2 | 59 | 2 | 8 | 19 | 8 | 0 | 0 | 0 | 0 | 3 | 4 | 9 | 8 | 6 | 12 | 7 | 64 |
| | BJ-76 | 4 | 3 | 0 | 0 | 1 | 11 | 0 | 1 | 18 | 0 | 0 | 1 | 60 | 3 | 5 | 11 | 4 | 0 | 0 | 0 | 0 | 3 | 5 | 14 | 13 | 9 | 11 | 4 | 39 |
| | BTM-1 | 0 | 0 | 0 | 0 | 0 | 1 | 0 | 0 | 1 | 0 | 0 | 0 | 5 | 0 | 1 | 1 | 0 | 0 | 0 | 0 | 0 | 1 | 0 | 1 | 1 | 1 | 1 | 0 | 1 |
| | BTM-8 | 0 | 0 | 0 | 0 | 0 | 0 | 0 | 0 | 1 | 0 | 0 | 0 | 2 | 0 | 1 | 0 | 0 | 0 | 0 | -1 | 0 | 0 | 0 | 1 | 0 | 0 | 0 | 0 | 1 |
| | F-1D7 | 0 | 0 | 0 | 0 | 0 | 0 | 0 | 0 | 0 | 0 | 0 | 0 | 1 | 0 | 0 | 0 | 0 | 0 | 0 | 0 | 0 | 0 | 0 | 0 | 0 | 0 | 0 | 0 | 0 |
| | F-1E7 | 0 | 0 | 0 | 0 | 0 | 0 | 0 | 0 | 0 | 0 | 0 | 0 | 1 | 0 | 0 | 0 | 0 | 0 | 0 | 0 | 0 | 0 | 0 | 0 | 0 | 0 | 0 | 0 | 0 |
| | F-2B4 | 0 | 0 | 0 | 0 | 0 | 0 | 0 | 0 | 0 | 0 | 0 | 0 | 0 | 0 | 0 | 0 | 0 | 0 | 0 | -1 | 0 | 0 | 0 | 0 | 0 | 0 | 0 | 0 | 0 |
| | F-3E2 | 0 | 0 | 0 | 0 | 0 | 0 | 0 | 0 | 0 | 0 | 0 | 0 | 0 | 0 | 0 | 0 | 0 | 0 | 0 | 0 | 0 | 0 | 0 | 0 | 0 | 0 | 0 | 0 | 0 |
| | FDX-01 | 3 | 2 | 0 | 0 | 0 | 7 | 0 | 1 | 14 | 0 | 0 | 2 | 65 | 2 | 6 | 10 | 4 | 0 | 0 | 0 | 0 | 2 | 4 | 15 | 11 | 7 | 11 | 4 | 40 |
| | FIND28 | 1 | 0 | 0 | 0 | 0 | 1 | 0 | 0 | 3 | 0 | 0 | 1 | 15 | 0 | 1 | 2 | 1 | 0 | 0 | 0 | 0 | 0 | 1 | 4 | 3 | 2 | 3 | 1 | 9 |
| | MCD022 Fab | 1 | 1 | 0 | 0 | 0 | 2 | 0 | 0 | 2 | 0 | 0 | 1 | 16 | 0 | 0 | 0 | 0 | 0 | 0 | 0 | 0 | 0 | 0 | 3 | 3 | 3 | 3 | 5 | 4 |
| | MCD022 Fab2 | 1 | 1 | 0 | 0 | 0 | 3 | 0 | 0 | 3 | 0 | 0 | 1 | 22 | 0 | 1 | 1 | 0 | 0 | 0 | 0 | 0 | 0 | 0 | 4 | 2 | 3 | 4 | 4 | 9 |
| | MCD024 Fab | 3 | 2 | 0 | 0 | 0 | 5 | 0 | 0 | 6 | 0 | 0 | 1 | 35 | 0 | 0 | 6 | 2 | 0 | 0 | 0 | 0 | 1 | 2 | 7 | 5 | 5 | 3 | 4 | 14 |
| | MCD024 Fab2 | 2 | 1 | 0 | 0 | 0 | 4 | 0 | 0 | 5 | 0 | 0 | 1 | 30 | 0 | 0 | 5 | 2 | 0 | 0 | 0 | 0 | 1 | 2 | 7 | 5 | 5 | 3 | 4 | 12 |
| | S4-20 | 4 | 2 | 0 | 0 | 0 | 6 | 0 | 1 | 6 | 0 | 0 | 1 | 100 | 4 | 8 | 17 | 7 | 0 | 0 | 0 | 0 | 6 | 9 | 8 | 5 | 5 | 3 | 2 | 54 |

### Table S4. Lineage 2 S / N ratio vs RAP

| Capture antibody | Detector antibody | | | | | | | | | | | | | | | | | | | | | | | | | | | | |
|---|---|---|---|---|---|---|---|---|---|---|---|---|---|---|---|---|---|---|---|---|---|---|---|---|---|---|---|---|---|
| | 11H2/ 11K1 | 15H3/ 15K3 | 16H2/ 16K1 | 17H2/ 17K3 | 18H2/ 18K2 | 1E7 | 20H3/ 20K2 | 52H3/ 52K2 | 5E3 | 79H2/ 79K2 | 7H3/ 7K3 | 90H3/ 90K3 | A194 -01 | BJ -03 | BJ -76 | BTM -1 | BTM -8 | F-1 D7 | F-1 E7 | F -2B4 | F-3 E2 | FDX -01 | FIND 28 | KI24 | MCD 022 Fab | MCD 022 Fab2 | MCD 024 Fab | MCD 024 Fab2 | S4-20 |
| 11H2/11K1 | 1 | 0 | 0 | 0 | 0 | 3 | 0 | 0 | 2 | 0 | 0 | 0 | 3 | 1 | 16 | 5 | 2 | 1 | 2 | 0 | 1 | 14 | 2 | 4 | 2 | 2 | 5 | 2 | 19 |
| 15H3/15K3 | 0 | 0 | 0 | 0 | 0 | 0 | 0 | 0 | 0 | 0 | 0 | 0 | 0 | 0 | 2 | 0 | 0 | 0 | 0 | 0 | 0 | 1 | 0 | 0 | 0 | 0 | 0 | 0 | 1 |
| 16H2/16K1 | 1 | 1 | 0 | 0 | 0 | 3 | 0 | 1 | 6 | 0 | 0 | 1 | 10 | 1 | 12 | 7 | 3 | 1 | 1 | 0 | 1 | 13 | 1 | 6 | 5 | 6 | 9 | 4 | 14 |
| 17H2/17K3 | 0 | 0 | 0 | 0 | 0 | 0 | 0 | 0 | 0 | 0 | 0 | 0 | 0 | 0 | 0 | 0 | 0 | 0 | 0 | 0 | 0 | 0 | 0 | 1 | 1 | 1 | 1 | 1 | 0 |
| 18H2/18K2 | 1 | 0 | 0 | 0 | 0 | 3 | 0 | 0 | 3 | 0 | 0 | 1 | 1 | 0 | 0 | 4 | 2 | 0 | 0 | 0 | 0 | 0 | 1 | 5 | 2 | 3 | 5 | 6 | 15 |
| 1E7 | 4 | 2 | 0 | 1 | 1 | 5 | 0 | 0 | 3 | 0 | 0 | 0 | 5 | 2 | 27 | 22 | 11 | 2 | 7 | 0 | 2 | 41 | 6 | 16 | 3 | 6 | 16 | 11 | 75 |
| 20H3/20K2 | 0 | 0 | 0 | 0 | 0 | 1 | 0 | 0 | 1 | 0 | 0 | 0 | 2 | 0 | 0 | 1 | 1 | 0 | 0 | 0 | 0 | - | 0 | 1 | 1 | 1 | 2 | 3 | 4 |
| 52H3/52K2 | 0 | 0 | 0 | 0 | 0 | 0 | 0 | 0 | 0 | 0 | 0 | 0 | 1 | 0 | 0 | 1 | 0 | 0 | 0 | 0 | 0 | 0 | 0 | 1 | 1 | 0 | 1 | 3 | 1 |
| 5E3 | 2 | 1 | 0 | 1 | 1 | 2 | 1 | 0 | 1 | 0 | 0 | 0 | 2 | 2 | 48 | 32 | 16 | 3 | 9 | 1 | 2 | 42 | 10 | 22 | 2 | 2 | 22 | 17 | 100 |
| 79H2/79K2 | 0 | 0 | 0 | 0 | 0 | 0 | 0 | 0 | 0 | 0 | 0 | 0 | 0 | 0 | 0 | 0 | 0 | 0 | 0 | 0 | 0 | 0 | 0 | 0 | 0 | 0 | 0 | 5 | 1 |
| 7H3/7K3 | 0 | 0 | 0 | 0 | 0 | 3 | 0 | 0 | 2 | 0 | 0 | 0 | 14 | 1 | 7 | 2 | 1 | 1 | 1 | 0 | 0 | 8 | 2 | 2 | 2 | 2 | 2 | 1 | 25 |
| 90H3/90K3 | 0 | 0 | 0 | 0 | 0 | 1 | 0 | 0 | 0 | 0 | 0 | 0 | 1 | 0 | 0 | 1 | 0 | 0 | 0 | 0 | 0 | 0 | 1 | 1 | 1 | 1 | 1 | 12 | 2 |
| A194-01 | 1 | 0 | 0 | 0 | 0 | 1 | 0 | 0 | 1 | 0 | 0 | 0 | 1 | 2 | 6 | 7 | 3 | 0 | 0 | 0 | 0 | 3 | 3 | 6 | 1 | 1 | 6 | 3 | 24 |
| BJ-03 | 13 | 8 | 0 | 1 | 2 | 23 | 1 | 3 | 31 | 0 | 0 | 4 | 60 | 4 | 15 | 35 | 16 | 0 | 0 | 0 | 0 | 7 | 8 | 18 | 12 | 13 | 29 | 16 | 130 |
| BJ-76 | 10 | 7 | 0 | 1 | 1 | 25 | 1 | 2 | 29 | 0 | 0 | 3 | 53 | 4 | 7 | 20 | 8 | 0 | 0 | 0 | 0 | 8 | 11 | 30 | 26 | 21 | 34 | 12 | 126 |
| BTM-1 | 1 | 1 | 0 | 0 | 0 | 2 | 0 | 0 | 2 | 0 | 0 | 0 | 4 | 0 | 2 | 1 | 1 | 0 | 0 | 0 | 0 | 1 | 1 | 2 | 1 | 1 | 1 | 1 | 1 |
| BTM-8 | 1 | 1 | 0 | 0 | 0 | 1 | 0 | 0 | 1 | 0 | 0 | 0 | 3 | 0 | 2 | 1 | 1 | 0 | 0 | 0 | 0 | 1 | 1 | 2 | 1 | 1 | 1 | 1 | 3 |
| F-1D7 | 0 | 0 | 0 | 0 | 0 | 0 | 0 | 0 | 0 | 0 | 0 | 0 | 1 | 0 | 0 | 0 | 0 | 0 | 0 | 0 | 0 | 0 | 0 | 0 | 0 | 0 | 0 | 0 | 1 |
| F-1E7 | 0 | 0 | 0 | 0 | 0 | 0 | 0 | 0 | 0 | 0 | 0 | 0 | 1 | 0 | 0 | 0 | 0 | 0 | 0 | 0 | 0 | 0 | 0 | 0 | 1 | 0 | 0 | 0 | 1 |
| F-2B4 | 0 | 0 | 0 | 0 | 0 | 0 | 0 | 0 | 0 | 0 | 0 | 0 | 0 | 0 | 0 | 0 | 1 | 0 | 0 | 0 | 0 | 0 | 0 | 0 | 0 | 0 | 0 | 0 | 0 |
| F-3E2 | 0 | 0 | 0 | 0 | 0 | 0 | 0 | 0 | 0 | 0 | 0 | 0 | 0 | 0 | 0 | 0 | 0 | 0 | 0 | 0 | 0 | 0 | 0 | 0 | 0 | 0 | 0 | 0 | 0 |
| FDX-01 | 8 | 5 | 0 | 1 | 1 | 20 | 1 | 2 | 29 | 1 | 0 | 5 | 29 | 3 | 16 | 27 | 11 | 0 | 0 | 0 | 0 | 6 | 9 | 23 | 15 | 15 | 25 | 12 | 105 |
| FIND28 | 2 | 1 | 0 | 0 | 0 | 4 | 0 | 1 | 6 | 0 | 0 | 1 | 15 | 1 | 3 | 6 | 2 | 0 | 0 | 0 | 0 | 1 | 2 | 5 | 7 | 4 | 10 | 3 | 22 |
| MCD022 Fab | 1 | 0 | 0 | 0 | 0 | 1 | 0 | 0 | 1 | 0 | 0 | 0 | 1 | 0 | 0 | 1 | 0 | 0 | 0 | 0 | 0 | 0 | 0 | 5 | 1 | 1 | 6 | 13 | 1 |
| MCD022 Fab2 | 1 | 0 | 0 | 0 | 0 | 1 | 0 | 0 | 1 | 0 | 0 | 0 | 1 | 0 | 1 | 1 | 1 | 0 | 0 | 0 | 0 | 1 | 1 | 7 | 0 | 0 | 1 | 4 | 2 |
| MCD024 Fab | 7 | 4 | 0 | 0 | 1 | 12 | 0 | 1 | 13 | 0 | 0 | 3 | 54 | 1 | 1 | 15 | 5 | 0 | 0 | 0 | 0 | 1 | 5 | 22 | 10 | 11 | 7 | 11 | 38 |
| MCD024 Fab2 | 6 | 3 | 0 | 0 | 1 | 11 | 0 | 1 | 13 | 0 | 0 | 2 | 58 | 1 | 1 | 12 | 4 | 0 | 0 | 0 | 0 | 1 | 5 | 15 | 8 | 7 | 6 | 3 | 25 |
| S4-20 | 9 | 5 | 0 | 1 | 1 | 13 | 0 | 1 | 13 | 0 | 0 | 2 | 100 | 7 | 12 | 26 | 13 | 0 | 0 | 0 | 0 | 16 | 19 | 19 | 6 | 10 | 6 | 4 | 95 |

### Table S5. Lineage 3 S - N ratio vs RAP

| Capture antibody | Detector antibody: 11H2/11K1 | 15H3/15K3 | 16H2/16K1 | 17H2/17K3 | 18H2/18K2 | 1E7 | 20H3/20K2 | 52H3/52K2 | 5E3 | 79H2/79K2 | 7H3/7K3 | 90H3/90K3 | A194-01 | BJ-03 | BJ-76 | BTM-1 | BTM-8 | F-1D7 | F-1E7 | F-2B4 | F-3E2 | FDX-01 | FIND28 | KI24 | MCD022 Fab | MCD022 Fab2 | MCD024 Fab | MCD024 Fab2 | S4-20 |
|---|---|---|---|---|---|---|---|---|---|---|---|---|---|---|---|---|---|---|---|---|---|---|---|---|---|---|---|---|---|
| 11H2/11K1 | 1 | 0 | 0 | 0 | 0 | 3 | 0 | 1 | 3 | 0 | 0 | 2 | 30 | 0 | 2 | 2 | 1 | 0 | 0 | 0 | 0 | 1 | 1 | 3 | 5 | 4 | 3 | 1 | 10 |
| 15H3/15K3 | 2 | 0 | 0 | 0 | 0 | 0 | 0 | 0 | 0 | 0 | 0 | 0 | 1 | 0 | 0 | 0 | 0 | 0 | 0 | 0 | 0 | 0 | 0 | 0 | 0 | 0 | 0 | 0 | 0 |
| 16H2/16K1 | 4 | 0 | 0 | 0 | 0 | 1 | 0 | 1 | 3 | 0 | 0 | 1 | 23 | 1 | 3 | 1 | 0 | 0 | 0 | 0 | 0 | 1 | 1 | 2 | 5 | 4 | 4 | 1 | 14 |
| 17H2/17K3 | 0 | 0 | 0 | 0 | 0 | 0 | 0 | 0 | 0 | 0 | 0 | 0 | 1 | 0 | 0 | 0 | 0 | 0 | 0 | 0 | 0 | 0 | 0 | 0 | 1 | 0 | 0 | -1 | 1 |
| 18H2/18K2 | 0 | 1 | 0 | 0 | 0 | 3 | 0 | 1 | 3 | 0 | 0 | 2 | 43 | 0 | 2 | 2 | 1 | 0 | 0 | 0 | 0 | 1 | 1 | 3 | 4 | 4 | 2 | -1 | 10 |
| 1E7 | 6 | 4 | 0 | 0 | 0 | 8 | 0 | 3 | 12 | 1 | 0 | 6 | 157 | 2 | 16 | 10 | 3 | 0 | 0 | 0 | 0 | 5 | 2 | 7 | 15 | 19 | 12 | 7 | 69 |
| 20H3/20K2 | 0 | 0 | 0 | 0 | 0 | 0 | 0 | 0 | 1 | 0 | 0 | 0 | 4 | 0 | 0 | 0 | 0 | 0 | 0 | 0 | 0 | - | 0 | 1 | 1 | 1 | 1 | -2 | 2 |
| 52H3/52K2 | 0 | 0 | 0 | 0 | 0 | 1 | 0 | 0 | 1 | 0 | 0 | 1 | 11 | 0 | 0 | 0 | 0 | 0 | 0 | 0 | 0 | 0 | 0 | 1 | 2 | 1 | 1 | -1 | 2 |
| 5E3 | 16 | 9 | 0 | 1 | 2 | 17 | 1 | 5 | 18 | 1 | 0 | 7 | 224 | 5 | 24 | 15 | 5 | 0 | 0 | 0 | 0 | 14 | 4 | 13 | 20 | 23 | 16 | 11 | 99 |
| 79H2/79K2 | 0 | 0 | 0 | 0 | 0 | 0 | 0 | 0 | 0 | 0 | 0 | 0 | 4 | 0 | 0 | 0 | 0 | 0 | 0 | 0 | 0 | 0 | 0 | 0 | 0 | 0 | 0 | -2 | 0 |
| 7H3/7K3 | 5 | 0 | 0 | 0 | 0 | 0 | 0 | 0 | 1 | 0 | 0 | 0 | 16 | 0 | 0 | 0 | 0 | 0 | 0 | 0 | 0 | 0 | 0 | 0 | 1 | 1 | 1 | 0 | 2 |
| 90H3/90K3 | 0 | 0 | 0 | 0 | 0 | 1 | 0 | 0 | 1 | 0 | 0 | 1 | 12 | 0 | 0 | 1 | 0 | 0 | 0 | 0 | 0 | 0 | 0 | 1 | 1 | 1 | 1 | -3 | 2 |
| A194-01 | 2 | 2 | 0 | 0 | 0 | 6 | 0 | 2 | 5 | 0 | 0 | 3 | 70 | 1 | 3 | 5 | 2 | 0 | 0 | 0 | 0 | 1 | 2 | 7 | 7 | 8 | 7 | 3 | 24 |
| BJ-03 | 2 | 2 | 0 | 0 | 0 | 4 | 0 | 1 | 8 | 0 | 0 | 2 | 48 | 1 | 3 | 3 | 1 | 0 | 0 | 1 | 0 | 1 | 1 | 4 | 6 | 4 | 8 | 5 | 36 |
| BJ-76 | 2 | 2 | 0 | 0 | 0 | 6 | 0 | 1 | 10 | 0 | 0 | 2 | 48 | 1 | 2 | 2 | 1 | 0 | -1 | 0 | 0 | 2 | 2 | 9 | 11 | 8 | 9 | 4 | 32 |
| BTM-1 | 3 | 0 | 0 | 0 | 0 | 0 | 0 | 0 | 0 | 0 | 0 | 0 | 3 | 0 | 0 | 0 | 0 | 0 | 0 | 0 | 0 | 0 | 0 | 0 | 0 | 0 | 0 | 0 | 3 |
| BTM-8 | 0 | 0 | 0 | 0 | 0 | 0 | 0 | 0 | 0 | 0 | 0 | 0 | 1 | 0 | 0 | 0 | 0 | 0 | 0 | -1 | 0 | 0 | 0 | 0 | 0 | 0 | 0 | 0 | 1 |
| F-1D7 | 0 | 0 | 0 | 0 | 0 | 0 | 0 | 0 | 0 | 0 | 0 | 0 | 0 | 0 | 0 | 0 | 0 | 0 | 0 | 0 | 0 | 0 | 0 | 0 | 0 | 0 | 0 | 0 | 0 |
| F-1E7 | 0 | 0 | 0 | 0 | 0 | 0 | 0 | 0 | 0 | 0 | 0 | 0 | 0 | 0 | 0 | 0 | 0 | 0 | -1 | 0 | 0 | 0 | 0 | 0 | 0 | 0 | 0 | 0 | 0 |
| F-2B4 | 0 | 0 | 0 | 0 | 0 | 0 | 0 | 0 | 0 | 0 | 0 | 0 | 0 | 0 | 0 | 0 | 0 | 0 | -2 | -13 | 0 | 0 | 0 | 0 | 0 | 0 | 0 | 0 | 0 |
| F-3E2 | 0 | 0 | 0 | 0 | 0 | 0 | 0 | 0 | 0 | 0 | 0 | 0 | 0 | 0 | 0 | 0 | 0 | 0 | 0 | 0 | 0 | 0 | 0 | 0 | 0 | 0 | 0 | 0 | 0 |
| FDX-01 | 2 | 2 | 0 | 0 | 0 | 8 | 0 | 3 | 7 | 1 | 0 | 7 | 121 | 1 | 3 | 2 | 1 | 0 | 0 | 0 | 0 | 1 | 2 | 10 | 18 | 13 | 7 | 3 | 28 |
| FIND28 | 1 | 0 | 0 | 0 | 0 | 2 | 0 | 1 | 2 | 0 | 0 | 2 | 25 | 0 | 1 | 0 | 0 | 0 | -1 | 0 | 0 | 0 | 1 | 3 | 5 | 3 | 2 | 1 | 5 |
| MCD022 Fab | 2 | 2 | 0 | 0 | 0 | 8 | 0 | 3 | 5 | 0 | 0 | 5 | 112 | 0 | 2 | 2 | 1 | 0 | 0 | 0 | 0 | 1 | 2 | 7 | 8 | 11 | 6 | 1 | 16 |
| MCD022 Fab2 | 2 | 2 | 0 | 0 | 0 | 5 | 0 | 1 | 6 | 0 | 0 | 1 | 63 | 1 | 4 | 4 | 1 | 0 | 0 | 0 | 0 | 2 | 1 | 6 | 4 | 8 | 8 | 2 | 36 |
| MCD024 Fab | 3 | 2 | 0 | 0 | 0 | 6 | 0 | 1 | 9 | 0 | 0 | 3 | 71 | 1 | 3 | 4 | 1 | 0 | 0 | 0 | 0 | 2 | 2 | 9 | 8 | 10 | 5 | 0 | 25 |
| MCD024 Fab2 | 3 | 2 | 0 | 0 | 0 | 6 | 0 | 1 | 7 | 0 | 0 | 3 | 64 | 1 | 2 | 4 | 1 | 0 | 0 | 0 | 0 | 2 | 2 | 8 | 9 | 10 | 4 | 1 | 22 |
| S4-20 | 4 | 2 | 0 | 0 | 0 | 6 | 0 | 1 | 8 | 0 | 0 | 3 | 100 | 2 | 4 | 3 | 1 | 0 | 0 | 0 | 0 | 3 | 4 | 8 | 9 | 9 | 4 | 2 | 36 |

### Table S6. Lineage 3 S / N ratio vs RAP

| | | Detector antibody | | | | | | | | | | | | | | | | | | | | | | | | | | | | |
|---|---|---|---|---|---|---|---|---|---|---|---|---|---|---|---|---|---|---|---|---|---|---|---|---|---|---|---|---|---|---|
| | | 11H2/11K1 | 15H3/15K3 | 16H2/16K1 | 17H2/17K3 | 18H2/18K2 | 1E7 | 20H3/20K2 | 52H3/52K2 | 5E3 | 79H2/79K2 | 7H3/7K3 | 90H3/90K3 | A194-01 | BJ-03 | BJ-76 | BTM-1 | BTM-8 | F-1D7 | F-1E7 | F-2B4 | F-3E2 | FDX-01 | FIND 28 | KI24 | MCD 022 Fab | MCD 022 Fab2 | MCD 024 Fab | MCD 024 Fab2 | S4-20 |
| Capture antibody | 11H2/11K1 | 3 | 1 | 1 | 1 | 1 | 14 | 1 | 4 | 6 | 2 | 1 | 4 | 11 | 2 | 5 | 7 | 3 | 1 | 1 | 1 | 1 | 5 | 5 | 12 | 10 | 10 | 14 | 8 | 41 |
| | 15H3/15K3 | 7 | 1 | 1 | 1 | 1 | 1 | 1 | 1 | 1 | 1 | 1 | 1 | 2 | 1 | 2 | 1 | 1 | 1 | 1 | 1 | 1 | 1 | 1 | 1 | 2 | 1 | 2 | 1 | 2 |
| | 16H2/16K1 | 13 | 2 | 1 | 1 | 1 | 7 | 1 | 4 | 13 | 2 | 1 | 5 | 27 | 3 | 9 | 5 | 3 | 1 | 1 | 1 | 1 | 5 | 3 | 8 | 18 | 17 | 23 | 8 | 51 |
| | 17H2/17K3 | 1 | 1 | 1 | 1 | 1 | 2 | 1 | 1 | 2 | 1 | 1 | 1 | 3 | 1 | 2 | 2 | 1 | 1 | 1 | 1 | 1 | 2 | 2 | 2 | 3 | 3 | 3 | 0 | 5 |
| | 18H2/18K2 | 2 | 1 | 1 | 1 | 1 | 10 | 1 | 4 | 8 | 2 | 1 | 4 | 22 | 2 | 6 | 6 | 3 | 1 | 1 | 1 | 1 | 5 | 4 | 12 | 4 | 9 | 10 | 1 | 33 |
| | 1E7 | 8 | 4 | 1 | 2 | 3 | 12 | 2 | 2 | 8 | 2 | 1 | 2 | 20 | 6 | 36 | 26 | 11 | 1 | 1 | 1 | 1 | 19 | 8 | 26 | 12 | 22 | 54 | 28 | 207 |
| | 20H3/20K2 | 1 | 1 | 1 | 1 | 1 | 3 | 1 | 2 | 3 | 1 | 1 | 2 | 7 | 2 | 2 | 2 | 2 | 1 | 1 | 1 | 1 | - | 2 | 4 | 2 | 5 | 4 | 0 | 9 |
| | 52H3/52K2 | 2 | 2 | 1 | 1 | 1 | 3 | 2 | 2 | 2 | 1 | 1 | 2 | 5 | 2 | 2 | 3 | 2 | 2 | 2 | 1 | 2 | 2 | 2 | 4 | 3 | 3 | 4 | 0 | 7 |
| | 5E3 | 7 | 4 | 2 | 3 | 4 | 7 | 3 | 2 | 4 | 2 | 2 | 2 | 10 | 14 | 65 | 41 | 19 | 1 | 1 | 1 | 1 | 49 | 13 | 35 | 7 | 8 | 69 | 52 | 279 |
| | 79H2/79K2 | 1 | 1 | 1 | 1 | 1 | 3 | 1 | 1 | 1 | 1 | 1 | 2 | 3 | 1 | 1 | 1 | 1 | 1 | 1 | 1 | 1 | 1 | 1 | 2 | 1 | 2 | 2 | 0 | 3 |
| | 7H3/7K3 | | | | | | | | | | | | | | | | | | | | | | | | | | | | | |
| | 90H3/90K3 | 16 | 1 | 1 | 1 | 1 | 2 | 1 | 2 | 3 | 1 | 1 | 2 | 28 | 1 | 2 | 2 | 1 | 1 | 1 | 1 | 1 | 2 | 2 | 3 | 5 | 4 | 5 | 2 | 7 |
| | A194-01 | 2 | 2 | 1 | 1 | 1 | 3 | 1 | 2 | 2 | 1 | 1 | 2 | 5 | 2 | 2 | 3 | 2 | 1 | 1 | 1 | 1 | 2 | 3 | 6 | 2 | 3 | 4 | 0 | 11 |
| | BJ-03 | 3 | 2 | 1 | 2 | 2 | 4 | 2 | 2 | 3 | 1 | 1 | 1 | 4 | 4 | 12 | 23 | 9 | 2 | 1 | 1 | 1 | 7 | 8 | 16 | 3 | 6 | 35 | 17 | 91 |
| | BJ-76 | 8 | 6 | 1 | 1 | 2 | 16 | 1 | 4 | 25 | 1 | 1 | 6 | 47 | 4 | 9 | 10 | 5 | 1 | 1 | 1 | 1 | 5 | 4 | 14 | 18 | 15 | 35 | 26 | 117 |
| | BTM-1 | 9 | 7 | 1 | 1 | 2 | 25 | 1 | 4 | 28 | 1 | 1 | 6 | 52 | 4 | 8 | 8 | 3 | 1 | 0 | 1 | 1 | 8 | 8 | 34 | 36 | 30 | 43 | 18 | 150 |
| | BTM-8 | 10 | 1 | 1 | 1 | 1 | 2 | 1 | 1 | 2 | 1 | 1 | 2 | 6 | 1 | 1 | 1 | 1 | 1 | 1 | 1 | 1 | 1 | 1 | 2 | 2 | 2 | 3 | 2 | 12 |
| | F-1D7 | 2 | 1 | 1 | 1 | 1 | 1 | 1 | 1 | 1 | 1 | 1 | 1 | 4 | 1 | 2 | 1 | 1 | 1 | 1 | 1 | 1 | 2 | 1 | 1 | 1 | 2 | 2 | 1 | 3 |
| | F-1E7 | 1 | 1 | 1 | 1 | 1 | 2 | 1 | 1 | 1 | 1 | 1 | 1 | 1 | 1 | 1 | 1 | 1 | 1 | 1 | 1 | 1 | 1 | 1 | 1 | 1 | 1 | 1 | 1 | 1 |
| | F-2B4 | 1 | 1 | 1 | 1 | 1 | 2 | 1 | 1 | 1 | 1 | 1 | 1 | 1 | 1 | 1 | 1 | 1 | 1 | 0 | 1 | 1 | 1 | 1 | 1 | 1 | 1 | 1 | 1 | 1 |
| | F-3E2 | 1 | 1 | 1 | 1 | 1 | 2 | 1 | 2 | 1 | 1 | 2 | 1 | 1 | 1 | 1 | 1 | 1 | 1 | 0 | 1 | 1 | 1 | 1 | 1 | 1 | 1 | 1 | 1 | 1 |
| | FDX-01 | 1 | 1 | 1 | 1 | 1 | 1 | 1 | 1 | 1 | 1 | 1 | 1 | 1 | 1 | 1 | 1 | 1 | 1 | 1 | 1 | 1 | 1 | 1 | 1 | 1 | 1 | 1 | 1 | 1 |
| | FIND28 | 8 | 6 | 1 | 1 | 2 | 29 | 2 | 9 | 23 | 3 | 1 | 20 | 70 | 4 | 13 | 7 | 4 | 1 | 1 | 1 | 1 | 6 | 6 | 21 | 44 | 29 | 26 | 17 | 104 |
| | MCD022 Fab | 4 | 3 | 1 | 2 | 1 | 10 | 2 | 6 | 6 | 3 | 2 | 11 | 39 | 2 | 4 | 3 | 2 | 1 | 0 | 1 | 2 | 3 | 3 | 9 | 17 | 11 | 11 | 4 | 26 |
| | MCD022 Fab2 | 3 | 2 | 1 | 1 | 2 | 5 | 1 | 2 | 3 | 1 | 1 | 2 | 11 | 3 | 9 | 10 | 4 | 1 | 1 | 1 | 1 | 6 | 8 | 19 | 3 | 6 | 25 | 1 | 74 |
| | MCD024 Fab | 3 | 2 | 1 | 1 | 1 | 4 | 1 | 2 | 3 | 1 | 1 | 1 | 6 | 2 | 11 | 8 | 6 | 1 | 1 | 1 | 1 | 8 | 2 | 16 | 2 | 2 | 3 | 2 | 132 |
| | MCD024 Fab2 | 15 | 8 | 1 | 2 | 2 | 26 | 2 | 6 | 32 | 2 | 1 | 12 | 216 | 7 | 13 | 15 | 6 | 1 | 1 | 1 | 1 | 12 | 12 | 40 | 7 | 39 | 18 | 1 | 88 |
| | S4-20 | 16 | 9 | 1 | 2 | 3 | 36 | 2 | 6 | 44 | 2 | 1 | 13 | 186 | 4 | 9 | 19 | 6 | 1 | 1 | 1 | 1 | 10 | 9 | 32 | 20 | 25 | 17 | 2 | 93 |
| | | 17 | 9 | 1 | 2 | 2 | 27 | 2 | 6 | 28 | 2 | 1 | 11 | 100 | 7 | 16 | 10 | 4 | 1 | 1 | 1 | 1 | 14 | 13 | 26 | 24 | 29 | 15 | 9 | 95 |

### Table S7. Lineage 4 S - N ratio vs RAP

| | | Detector antibody | | | | | | | | | | | | | | | | | | | | | | | | | | | |
|---|---|---|---|---|---|---|---|---|---|---|---|---|---|---|---|---|---|---|---|---|---|---|---|---|---|---|---|---|---|
| | | 11H2/ 11K1 | 15H3/ 15K3 | 16H2/ 16K1 | 17H2/ 17K3 | 18H2/ 18K2 | 1E7 | 20H3/ 20K2 | 52H3/ 52K2 | 5E3 | 79H2/ 79K2 | 7H3/ 7K3 | 90H3/ 90K3 | A194 -01 | BJ -03 | BJ -76 | BTM -1 | BTM -8 | F-1 D7 | F-1 E7 | F -2B4 | F-3 E2 | FDX -01 | FIND 28 | KI24 | MCD 022 Fab | MCD 022 Fab2 | MCD 024 Fab | MCD 024 Fab2 | S4-20 |
| Capture antibody | 11H2/11K1 | 1 | 1 | 0 | 0 | 0 | 3 | 0 | 1 | 5 | 0 | 0 | 1 | 30 | 0 | 2 | 1 | 0 | 0 | 0 | 0 | 0 | 1 | 1 | 9 | 7 | 6 | 5 | 2 | 19 |
| | 15H3/15K3 | 0 | 0 | 0 | 0 | 0 | 0 | 0 | 0 | 0 | 0 | 0 | 0 | 1 | 0 | 0 | 0 | 0 | 0 | 0 | 0 | 0 | 0 | 0 | 0 | 0 | 0 | 0 | 0 | 0 |
| | 16H2/16K1 | 1 | 0 | 0 | 0 | 0 | 1 | 0 | 0 | 3 | 0 | 0 | 1 | 26 | 1 | 5 | 1 | 0 | 0 | 0 | 0 | 0 | 2 | 1 | 4 | 5 | 5 | 4 | 1 | 13 |
| | 17H2/17K3 | 0 | 0 | 0 | 0 | 0 | 0 | 0 | 0 | 1 | 0 | 0 | 0 | 2 | 0 | 0 | 0 | 0 | 0 | 0 | 0 | 0 | 0 | 0 | 1 | 1 | 1 | 1 | 0 | 2 |
| | 18H2/18K2 | 1 | 0 | 0 | 0 | 0 | 2 | 0 | 0 | 4 | 0 | 0 | 1 | 30 | 0 | 2 | 1 | 0 | 0 | 0 | 0 | 0 | 1 | 1 | 5 | 4 | 3 | 3 | 1 | 10 |
| | 1E7 | 9 | 5 | 0 | 0 | 1 | 9 | 0 | 2 | 13 | 0 | 0 | 4 | 104 | 2 | 14 | 5 | 2 | 0 | 0 | 0 | 0 | 6 | 3 | 16 | 16 | 15 | 12 | 7 | 69 |
| | 20H3/20K2 | 0 | 0 | 0 | 0 | 0 | 1 | 0 | 0 | 2 | 0 | 0 | 0 | 6 | 0 | 0 | 0 | 0 | 0 | 0 | 0 | 0 | - | 0 | 2 | 2 | 2 | 1 | 0 | 3 |
| | 52H3/52K2 | 0 | 0 | 0 | 0 | 0 | 0 | 0 | 0 | 0 | 0 | 0 | 0 | 3 | 0 | 0 | 0 | 0 | 0 | 0 | 0 | 0 | 0 | 0 | 0 | 1 | 0 | 0 | 0 | 1 |
| | 5E3 | 15 | 7 | 0 | 1 | 1 | 13 | 1 | 2 | 20 | 0 | 0 | 5 | 133 | 4 | 21 | 7 | 2 | 0 | 0 | 0 | 0 | 12 | 4 | 24 | 19 | 17 | 14 | 9 | 80 |
| | 79H2/79K2 | 0 | 0 | 0 | 0 | 0 | 0 | 0 | 0 | 0 | 0 | 0 | 0 | 1 | 0 | 0 | 0 | 0 | 0 | 0 | 0 | 0 | 0 | 0 | 0 | 0 | 0 | 0 | 0 | 0 |
| | 7H3/7K3 | 0 | 0 | 0 | 0 | 0 | 2 | 0 | 0 | 2 | 0 | 0 | 0 | 16 | 0 | 1 | 0 | 0 | 0 | 0 | 0 | 0 | 0 | 1 | 3 | 4 | 3 | 2 | 0 | 20 |
| | 90H3/90K3 | 0 | 0 | 0 | 0 | 0 | 0 | 0 | 0 | 1 | 0 | 0 | 0 | 7 | 0 | 0 | 0 | 0 | 0 | 0 | 0 | 0 | 0 | 0 | 1 | 1 | 1 | 1 | 0 | 2 |
| | A194-01 | 3 | 2 | 0 | 0 | 0 | 4 | 0 | 1 | 5 | 0 | 0 | 2 | 48 | 0 | 2 | 2 | 1 | 0 | 0 | -2 | 0 | 1 | 2 | 9 | 7 | 7 | 5 | 2 | 20 |
| | BJ-03 | 6 | 3 | 0 | 0 | 1 | 6 | 0 | 1 | 9 | 0 | 0 | 1 | 58 | 1 | 5 | 4 | 1 | 0 | 0 | -3 | 0 | 2 | 3 | 9 | 7 | 5 | 9 | 6 | 41 |
| | BJ-76 | 5 | 4 | 0 | 0 | 1 | 8 | 0 | 1 | 11 | 0 | 0 | 1 | 70 | 1 | 3 | 3 | 1 | 0 | 0 | 0 | 0 | 3 | 4 | 14 | 13 | 9 | 11 | 5 | 35 |
| | BTM-1 | 0 | 0 | 0 | 0 | 0 | 0 | 0 | 0 | 0 | 0 | 0 | 0 | 2 | 0 | 0 | 0 | 0 | 0 | 0 | 0 | 0 | 0 | 0 | 0 | 0 | 0 | 0 | 0 | 1 |
| | BTM-8 | 0 | 0 | 0 | 0 | 0 | 0 | 0 | 0 | 0 | 0 | 0 | 0 | 1 | 0 | 0 | 0 | 0 | 0 | 0 | 0 | 0 | 0 | 0 | 0 | 0 | 0 | 0 | 0 | 0 |
| | F-1D7 | 0 | 0 | 0 | 0 | 0 | 0 | 0 | 0 | 0 | 0 | 0 | 0 | 0 | 0 | 0 | 0 | 0 | 0 | 0 | 2 | 0 | 0 | 0 | 0 | 0 | 0 | 0 | 0 | 0 |
| | F-1E7 | 0 | 0 | 0 | 0 | 0 | 0 | 0 | 0 | 0 | 0 | 0 | 0 | 0 | 0 | 0 | 0 | 0 | 0 | 0 | 1 | 0 | 0 | 0 | 0 | 0 | 0 | 0 | 0 | 0 |
| | F-2B4 | 0 | 0 | 0 | 0 | 0 | 0 | 0 | 0 | 0 | 0 | 0 | 0 | 0 | 0 | 0 | 0 | 0 | 0 | 0 | -32 | 0 | 0 | 0 | 0 | 0 | 0 | 0 | 0 | 0 |
| | F-3E2 | 0 | 0 | 0 | 0 | 0 | 0 | 0 | 0 | 0 | 0 | 0 | 0 | 0 | 0 | 0 | 0 | 0 | 0 | 0 | 0 | 0 | 0 | 0 | 0 | 0 | 0 | 0 | 0 | 0 |
| | FDX-01 | 4 | 2 | 0 | 0 | 0 | 6 | 0 | 1 | 9 | 0 | 0 | 2 | 68 | 1 | 3 | 2 | 1 | 0 | 0 | 1 | 0 | 2 | 3 | 15 | 11 | 8 | 9 | 4 | 30 |
| | FIND28 | 1 | 1 | 0 | 0 | 0 | 1 | 0 | 0 | 3 | 0 | 0 | 0 | 19 | 0 | 1 | 1 | 0 | 0 | 0 | 0 | 0 | 0 | 1 | 5 | 4 | 2 | 3 | 1 | 8 |
| | MCD022 Fab | 3 | 2 | 0 | 0 | 0 | 5 | 0 | 1 | 6 | 0 | 0 | 2 | 43 | 0 | 1 | 1 | 0 | 0 | 0 | 0 | 0 | 1 | 1 | 8 | 7 | 7 | 6 | 2 | 12 |
| | MCD022 Fab2 | 3 | 1 | 0 | 0 | 0 | 5 | 0 | 1 | 6 | 0 | 0 | 2 | 55 | 1 | 2 | 2 | 1 | 0 | 0 | 0 | 0 | 1 | 1 | 9 | 8 | 8 | 7 | 3 | 25 |
| | MCD024 Fab | 5 | 3 | 0 | 0 | 0 | 7 | 0 | 1 | 10 | 0 | 0 | 1 | 60 | 1 | 3 | 2 | 1 | 0 | 0 | 0 | 0 | 3 | 3 | 15 | 10 | 10 | 7 | 3 | 28 |
| | MCD024 Fab2 | 5 | 3 | 0 | 0 | 0 | 6 | 0 | 1 | 8 | 0 | 0 | 1 | 57 | 1 | 3 | 2 | 1 | 0 | 0 | 0 | 0 | 2 | 3 | 13 | 9 | 10 | 6 | 3 | 26 |
| | S4-20 | 6 | 3 | 0 | 0 | 0 | 7 | 0 | 1 | 8 | 0 | 0 | 1 | 100 | 2 | 5 | 4 | 1 | 0 | 0 | 0 | 0 | 5 | 7 | 15 | 9 | 9 | 6 | 3 | 42 |

### Table S8. Lineage 4 S/N ratio vs RAP

| | | Detector antibody | | | | | | | | | | | | | | | | | | | | | | | | | | | |
|---|---|---|---|---|---|---|---|---|---|---|---|---|---|---|---|---|---|---|---|---|---|---|---|---|---|---|---|---|---|
| | | 11H2/ 11K1 | 15H3/ 15K3 | 16H2/ 16K1 | 17H2/ 17K3 | 18H2/ 18K2 | 1E7 | 20H3/ 20K2 | 52H3/ 52K2 | 5E3 | 79H2/ 79K2 | 7H3/ 7K3 | 90H3/ 90K3 | A194 -01 | BJ -03 | BJ -76 | BTM -1 | BTM -8 | F-1 D7 | F-1 E7 | F -2B4 | F-3 E2 | FDX -01 | FIND 28 | KI24 | MCD 022 Fab | MCD 022 Fab2 | MCD 024 Fab | MCD 024 Fab2 | S4-20 |
| Capture antibody | 11H2/11K1 | 5 | 1 | 1 | 1 | 1 | 10 | 1 | 2 | 9 | 1 | 1 | 2 | 18 | 2 | 7 | 5 | 2 | 0 | 0 | 1 | 0 | 6 | 7 | 18 | 11 | 15 | 20 | 10 | 75 |
| | 15H3/15K3 | 1 | 1 | 0 | 0 | 0 | 1 | 0 | 0 | 1 | 0 | 0 | 1 | 2 | 1 | 1 | 1 | 1 | 1 | 1 | 1 | 1 | 1 | 1 | 1 | 1 | 1 | 1 | 1 | 2 |
| | 16H2/16K1 | 3 | 2 | 0 | 1 | 1 | 4 | 1 | 2 | 13 | 1 | 0 | 3 | 63 | 3 | 16 | 5 | 2 | 0 | 1 | 1 | 0 | 9 | 4 | 16 | 18 | 23 | 15 | 7 | 52 |
| | 17H2/17K3 | 1 | 1 | 1 | 1 | 1 | 2 | 1 | 1 | 3 | 1 | 1 | 1 | 4 | 1 | 1 | 1 | 1 | 1 | 1 | 0 | 1 | 1 | 1 | 5 | 5 | 4 | 5 | 3 | 6 |
| | 18H2/18K2 | 3 | 1 | 1 | 1 | 1 | 8 | 1 | 1 | 12 | 1 | 1 | 2 | 17 | 2 | 7 | 3 | 2 | 1 | 1 | 0 | 0 | 5 | 3 | 16 | 11 | 9 | 12 | 8 | 38 |
| | 1E7 | 17 | 8 | 1 | 2 | 3 | 15 | 2 | 2 | 12 | 1 | 1 | 2 | 22 | 7 | 40 | 21 | 7 | 0 | 0 | 0 | 0 | 27 | 12 | 37 | 14 | 22 | 57 | 37 | 225 |
| | 20H3/20K2 | 2 | 1 | 1 | 1 | 1 | 3 | 0 | 1 | 5 | 1 | 0 | 1 | 15 | 1 | 2 | 2 | 1 | 0 | 1 | 0 | 1 | - | 2 | 8 | 9 | 8 | 9 | 3 | 17 |
| | 52H3/52K2 | 1 | 1 | 0 | 0 | 0 | 1 | 1 | 1 | 1 | 1 | 0 | 1 | 3 | 1 | 2 | 1 | 1 | 0 | 1 | 0 | 1 | 1 | 1 | 2 | 2 | 1 | 2 | 1 | 4 |
| | 5E3 | 11 | 6 | 1 | 2 | 3 | 10 | 3 | 1 | 5 | 1 | 1 | 1 | 10 | 13 | 62 | 24 | 9 | 0 | 1 | 0 | 1 | 49 | 18 | 35 | 8 | 8 | 60 | 46 | 310 |
| | 79H2/79K2 | 1 | 1 | 0 | 1 | 0 | 1 | 1 | 1 | 1 | 1 | 1 | 1 | 2 | 1 | 1 | 1 | 1 | 0 | 0 | 0 | 0 | 0 | 1 | 1 | 2 | 1 | 2 | 1 | 2 |
| | 7H3/7K3 | 2 | 1 | 0 | 1 | 1 | 8 | 1 | 1 | 11 | 1 | 1 | 2 | 21 | 1 | 4 | 2 | 1 | 0 | 0 | 0 | 0 | 2 | 4 | 11 | 15 | 17 | 12 | 3 | 92 |
| | 90H3/90K3 | 2 | 1 | 1 | 1 | 1 | 2 | 1 | 1 | 1 | 1 | 1 | 1 | 4 | 1 | 1 | 1 | 1 | 1 | 0 | 1 | 1 | 1 | 1 | 4 | 4 | 3 | 4 | 2 | 7 |
| | A194-01 | 4 | 2 | 1 | 1 | 1 | 3 | 1 | 1 | 2 | 1 | 1 | 1 | 4 | 2 | 8 | 7 | 3 | 0 | 0 | 0 | 0 | 5 | 6 | 13 | 3 | 5 | 26 | 10 | 75 |
| | BJ-03 | 21 | 13 | 0 | 2 | 3 | 24 | 1 | 4 | 33 | 1 | 1 | 6 | 81 | 5 | 16 | 13 | 5 | 0 | 0 | 0 | 0 | 8 | 12 | 35 | 22 | 21 | 48 | 33 | 161 |
| | BJ-76 | 23 | 15 | 0 | 2 | 3 | 37 | 1 | 4 | 36 | 1 | 1 | 5 | 80 | 5 | 10 | 9 | 3 | 0 | 0 | 0 | 0 | 15 | 17 | 64 | 53 | 41 | 60 | 28 | 149 |
| | BTM-1 | 1 | 1 | 0 | 1 | 1 | 1 | 1 | 1 | 1 | 1 | 1 | 1 | 7 | 1 | 1 | 1 | 1 | 0 | 0 | 0 | 0 | 1 | 1 | 2 | 2 | 2 | 2 | 1 | 5 |
| | BTM-8 | 1 | 1 | 0 | 0 | 0 | 1 | 0 | 1 | 1 | 0 | 0 | 0 | 3 | 1 | 1 | 1 | 0 | 1 | 1 | 0 | 0 | 1 | 1 | 1 | 1 | 1 | 1 | 1 | 2 |
| | F-1D7 | 0 | 0 | 0 | 0 | 0 | 1 | 0 | 0 | 0 | 0 | 0 | 1 | 1 | 0 | 0 | 0 | 0 | 0 | 0 | 1 | 0 | 0 | 0 | 1 | 1 | 0 | 1 | 1 | 1 |
| | F-1E7 | 1 | 1 | 0 | 1 | 1 | 1 | 1 | 1 | 0 | 1 | 1 | 1 | 1 | 1 | 1 | 1 | 1 | 1 | 0 | 1 | 1 | 0 | 1 | 1 | 1 | 1 | 1 | 1 | 1 |
| | F-2B4 | 0 | 1 | 1 | 1 | 1 | 1 | 0 | 0 | 1 | 0 | 0 | 0 | 1 | 1 | 0 | 1 | 0 | 1 | 0 | 0 | 1 | 1 | 0 | 2 | 2 | 2 | 1 | 1 | 1 |
| | F-3E2 | 0 | 0 | 0 | 0 | 0 | 1 | 1 | 0 | 1 | 0 | 0 | 0 | 1 | 0 | 0 | 0 | 1 | 0 | 0 | 0 | 0 | 1 | 1 | 1 | 1 | 1 | 1 | 1 | 0 |
| | FDX-01 | 16 | 7 | 0 | 2 | 2 | 19 | 1 | 4 | 29 | 1 | 1 | 5 | 41 | 3 | 11 | 7 | 2 | 0 | 0 | 1 | 0 | 7 | 10 | 49 | 37 | 29 | 55 | 22 | 189 |
| | FIND28 | 7 | 3 | 1 | 1 | 1 | 7 | 1 | 1 | 11 | 1 | 1 | 2 | 32 | 2 | 5 | 3 | 2 | 0 | 0 | 0 | 0 | 3 | 4 | 12 | 15 | 11 | 17 | 5 | 39 |
| | MCD022 Fab | 5 | 2 | 0 | 1 | 1 | 4 | 1 | 1 | 3 | 1 | 1 | 1 | 5 | 2 | 6 | 5 | 2 | 1 | 0 | 0 | 0 | 4 | 4 | 18 | 5 | 6 | 41 | 13 | 49 |
| | MCD022 Fab2 | 6 | 2 | 1 | 1 | 1 | 4 | 1 | 1 | 3 | 1 | 1 | 1 | 5 | 2 | 8 | 5 | 3 | 1 | 0 | 0 | 1 | 10 | 2 | 28 | 4 | 2 | 4 | 6 | 108 |
| | MCD024 Fab | 23 | 13 | 1 | 1 | 2 | 33 | 1 | 3 | 35 | 1 | 1 | 5 | 169 | 6 | 18 | 11 | 5 | 1 | 1 | 0 | 1 | 14 | 14 | 51 | 38 | 36 | 30 | 20 | 106 |
| | MCD024 Fab2 | 25 | 14 | 0 | 1 | 3 | 37 | 1 | 4 | 43 | 1 | 1 | 8 | 205 | 4 | 14 | 11 | 4 | 1 | 1 | 1 | 1 | 10 | 15 | 56 | 42 | 27 | 31 | 5 | 141 |
| | S4-20 | 27 | 14 | 1 | 2 | 3 | 30 | 2 | 4 | 44 | 1 | 1 | 7 | 100 | 5 | 15 | 12 | 4 | 0 | 0 | 0 | 0 | 28 | 30 | 55 | 27 | 34 | 24 | 16 | 154 |

### Table S9. Lineage 5:6 S - N ratio vs RAP

| Capture antibody | Detector antibody: 11H2/ 11K1 | 15H3/ 15K3 | 16H2/ 16K1 | 17H2/ 17K3 | 18H2/ 18K2 | 1E7 | 20H3/ 20K2 | 52H3/ 52K2 | 5E3 | 79H2/ 79K2 | 7H3/ 7K3 | 90H3/ 90K3 | A194 -01 | BJ -03 | BJ -76 | BTM -1 | BTM -8 | F-1 D7 | F-1 E7 | F -2B4 | F-3 E2 | FDX -01 | FIND 28 | KI24 | MCD 022 Fab | MCD 022 Fab2 | MCD 024 Fab | MCD 024 Fab2 | S4-20 |
|---|---|---|---|---|---|---|---|---|---|---|---|---|---|---|---|---|---|---|---|---|---|---|---|---|---|---|---|---|---|
| 11H2/11K1 | 0 | 0 | 0 | 0 | 0 | 3 | 0 | 0 | 5 | 0 | 0 | 1 | 13 | 0 | 0 | 8 | 3 | 0 | 0 | 0 | 0 | 0 | 1 | 4 | 3 | 2 | 9 | 4 | 38 |
| 15H3/15K3 | 0 | 0 | 0 | 0 | 0 | 0 | 0 | 0 | 0 | 0 | 0 | 0 | 0 | 0 | 0 | 0 | 0 | 0 | 0 | 0 | 0 | 0 | 0 | 0 | 0 | 0 | 0 | 0 | 1 |
| 16H2/16K1 | 0 | 0 | 0 | 0 | 0 | 2 | 0 | 0 | 6 | 0 | 0 | 1 | 20 | 0 | 0 | 10 | 3 | 0 | 0 | 0 | 0 | 0 | 1 | 3 | 3 | 3 | 11 | 4 | 42 |
| 17H2/17K3 | 0 | 0 | 0 | 0 | 0 | 0 | 0 | 0 | 0 | 0 | 0 | 0 | 0 | 0 | 0 | 0 | 0 | 0 | 0 | 0 | 0 | 0 | 0 | 0 | 0 | 0 | 1 | 0 | 0 |
| 18H2/18K2 | 1 | 0 | 0 | 0 | 0 | 3 | 0 | 0 | 4 | 0 | 0 | 1 | 2 | 0 | 0 | 9 | 3 | 0 | 0 | 0 | 0 | 0 | 1 | 4 | 2 | 2 | 8 | 4 | 31 |
| 1E7 | 4 | 3 | 0 | 0 | 1 | 10 | 0 | 1 | 16 | 0 | 0 | 3 | 73 | 0 | 1 | 46 | 15 | 0 | 0 | 0 | 0 | 0 | 4 | 11 | 10 | 10 | 35 | 19 | 174 |
| 20H3/20K2 | 0 | 0 | 0 | 0 | 0 | 1 | 0 | 0 | 1 | 0 | 0 | 0 | 3 | 0 | 0 | 2 | 1 | 0 | 0 | 0 | 0 | - | 0 | 1 | 1 | 0 | 3 | 1 | 8 |
| 52H3/52K2 | 0 | 0 | 0 | 0 | 0 | 0 | 0 | 0 | 0 | 0 | 0 | 0 | 3 | 0 | 0 | 0 | 0 | 0 | 0 | 0 | 0 | 0 | 0 | 1 | 0 | 0 | 1 | 0 | 2 |
| 5E3 | 8 | 6 | 0 | 0 | 1 | 15 | 0 | 2 | 23 | 0 | 0 | 3 | 94 | 0 | 1 | 55 | 18 | 0 | 0 | 0 | 0 | 0 | 7 | 20 | 12 | 12 | 39 | 23 | 191 |
| 79H2/79K2 | 0 | 0 | 0 | 0 | 0 | 0 | 0 | 0 | 0 | 0 | 0 | 0 | 1 | 0 | 0 | 0 | 0 | 0 | 0 | 0 | 0 | 0 | 0 | 0 | 0 | 0 | 0 | 0 | 0 |
| 7H3/7K3 | 0 | 0 | 0 | 0 | 0 | 1 | 0 | 0 | 2 | 0 | 0 | 0 | 7 | 0 | 0 | 2 | 0 | 0 | 0 | 0 | 0 | 0 | 1 | 1 | 1 | 1 | 5 | 1 | 30 |
| 90H3/90K3 | 0 | 0 | 0 | 0 | 0 | 1 | 0 | 0 | 1 | 0 | 0 | 0 | 4 | 0 | 0 | 1 | 0 | 0 | 0 | 0 | 0 | 0 | 0 | 1 | 1 | 0 | 1 | 0 | 4 |
| A194-01 | 1 | 1 | 0 | 0 | 0 | 5 | 0 | 1 | 5 | 0 | 0 | 1 | 24 | 1 | 4 | 9 | 3 | 0 | 0 | 0 | 0 | 1 | 2 | 7 | 3 | 3 | 11 | 4 | 41 |
| BJ-03 | 2 | 2 | 0 | 0 | 0 | 5 | 0 | 0 | 8 | 0 | 0 | 1 | 30 | 1 | 13 | 18 | 7 | 0 | 0 | 0 | 0 | 2 | 2 | 7 | 3 | 2 | 14 | 8 | 59 |
| BJ-76 | 5 | 5 | 0 | 0 | 1 | 18 | 0 | 1 | 23 | 0 | 0 | 1 | 56 | 3 | 29 | 38 | 14 | 0 | 0 | 0 | 0 | 6 | 9 | 27 | 10 | 7 | 43 | 23 | 208 |
| BTM-1 | 1 | 0 | 0 | 0 | 0 | 1 | 0 | 0 | 2 | 0 | 0 | 0 | 5 | 0 | 0 | 3 | 1 | 0 | 0 | 0 | 0 | 0 | 1 | 2 | 1 | 1 | 4 | 2 | 0 |
| BTM-8 | 0 | 0 | 0 | 0 | 0 | 1 | 0 | 0 | 1 | 0 | 0 | 0 | 3 | 0 | 0 | 2 | 1 | 0 | 0 | 0 | 0 | 0 | 0 | 1 | 1 | 0 | 3 | 1 | 11 |
| F-1D7 | 0 | 0 | 0 | 0 | 0 | 0 | 0 | 0 | 0 | 0 | 0 | 0 | 0 | 0 | 0 | 0 | 0 | 0 | 0 | 0 | 0 | 0 | 0 | 0 | 0 | 0 | 0 | 0 | 0 |
| F-1E7 | 0 | 0 | 0 | 0 | 0 | 0 | 0 | 0 | 0 | 0 | 0 | 0 | 0 | 0 | 0 | 0 | 0 | 0 | 0 | 0 | 0 | 0 | 0 | 0 | 0 | 0 | 0 | 0 | 1 |
| F-2B4 | 0 | 0 | 0 | 0 | 0 | 0 | 0 | 0 | 0 | 0 | 0 | 0 | 0 | 0 | 0 | 0 | 0 | 0 | 0 | -2 | 0 | 0 | 0 | 0 | 0 | 0 | 0 | 0 | 0 |
| F-3E2 | 0 | 0 | 0 | 0 | 0 | 0 | 0 | 0 | 0 | 0 | 0 | 0 | 0 | 0 | 0 | 0 | 0 | 0 | 0 | 0 | 0 | 0 | 0 | 0 | 0 | 0 | 0 | 0 | 0 |
| FDX-01 | 2 | 2 | 0 | 0 | 0 | 7 | 0 | 1 | 10 | 0 | 0 | 2 | 51 | 1 | 16 | 16 | 6 | 0 | 0 | 0 | 0 | 2 | 4 | 16 | 7 | 5 | 22 | 10 | 94 |
| FIND28 | 1 | 0 | 0 | 0 | 0 | 2 | 0 | 0 | 3 | 0 | 0 | 1 | 13 | 0 | 4 | 4 | 2 | 0 | 0 | 0 | 0 | 0 | 1 | 5 | 2 | 1 | 8 | 2 | 24 |
| MCD022 Fab | 1 | 1 | 0 | 0 | 0 | 5 | 0 | 1 | 5 | 0 | 0 | 2 | 6 | 0 | 0 | 0 | 0 | 0 | 0 | 0 | 0 | 0 | 0 | 8 | 4 | 4 | 11 | 4 | 1 |
| MCD022 Fab2 | 1 | 1 | 0 | 0 | 0 | 5 | 0 | 0 | 6 | 0 | 0 | 1 | 3 | 0 | 0 | 0 | 0 | 0 | 0 | 0 | 0 | 0 | 0 | 8 | 3 | 3 | 15 | 6 | 2 |
| MCD024 Fab | 7 | 4 | 0 | 0 | 1 | 16 | 0 | 1 | 16 | 0 | 0 | 2 | 55 | 0 | 0 | 42 | 16 | 0 | 0 | 0 | 0 | 0 | 5 | 24 | 9 | 7 | 29 | 17 | 133 |
| MCD024 Fab2 | 5 | 3 | 0 | 0 | 1 | 13 | 0 | 0 | 17 | 0 | 0 | 2 | 57 | 0 | 0 | 35 | 12 | 0 | 0 | 0 | 0 | 0 | 6 | 21 | 9 | 9 | 25 | 15 | 129 |
| S4-20 | 8 | 5 | 0 | 0 | 1 | 17 | 0 | 1 | 20 | 0 | 0 | 2 | 100 | 6 | 44 | 49 | 18 | 0 | 0 | 0 | 0 | 10 | 14 | 22 | 10 | 9 | 25 | 16 | 195 |

### Table S10. Lineage 5:6 S/N ratio v RAP

| | | Detector Antibody | | | | | | | | | | | | | | | | | | | | | | | | | | | | |
|---|---|---|---|---|---|---|---|---|---|---|---|---|---|---|---|---|---|---|---|---|---|---|---|---|---|---|---|---|---|---|
| | | 11H2/ 11K1 | 15H3/ 15K3 | 16H2/ 16K1 | 17H2/ 17K3 | 18H2/ 18K2 | 1E7 | 20H3/ 20K2 | 52H3/ 52K2 | 5E3 | 79H2/ 79K2 | 7H3/ 7K3 | 90H3/ 90K3 | A194 -01 | BJ -03 | BJ -76 | BTM -1 | BTM -8 | F-1 D7 | F-1 E7 | F -2B4 | F-3 E2 | FDX -01 | FIND 28 | KI24 | MCD 022 Fab | MCD 022 Fab2 | MCD 024 Fab | MCD 024 Fab2 | S4-20 |
| Capture antibody | 11H2/11K1 | 1 | 0 | 0 | 0 | 0 | 7 | 0 | 1 | 6 | 0 | 0 | 1 | 4 | 0 | 0 | 31 | 11 | 0 | 0 | 0 | 0 | 0 | 4 | 9 | 3 | 3 | 33 | 18 | 92 |
| | 15H3/15K3 | 0 | 0 | 0 | 0 | 0 | 0 | 0 | 0 | 1 | 0 | 0 | 0 | 0 | 0 | 0 | 1 | 0 | 0 | 0 | 0 | 0 | 0 | 0 | 1 | 0 | 0 | 2 | 1 | 4 |
| | 16H2/16K1 | 1 | 0 | 0 | 0 | 0 | 8 | 0 | 2 | 16 | 0 | 0 | 3 | 20 | 0 | 0 | 26 | 9 | 0 | 0 | 0 | 0 | 0 | 3 | 11 | 11 | 9 | 38 | 17 | 184 |
| | 17H2/17K3 | 0 | 0 | 0 | 0 | 0 | 1 | 0 | 0 | 1 | 0 | 0 | 0 | 0 | 0 | 0 | 0 | 0 | 0 | 0 | 0 | 0 | 0 | 0 | 1 | 1 | 1 | 7 | 2 | 0 |
| | 18H2/18K2 | 2 | 0 | 0 | 0 | 0 | 8 | 0 | 1 | 9 | 0 | 0 | 1 | 0 | 0 | 0 | 26 | 9 | 0 | 0 | 0 | 0 | 0 | 4 | 13 | 4 | 4 | 29 | 15 | 111 |
| | 1E7 | 5 | 2 | 0 | 1 | 2 | 11 | 1 | 1 | 8 | 0 | 0 | 1 | 8 | 0 | 1 | 145 | 42 | 0 | 0 | 0 | 0 | 0 | 11 | 32 | 6 | 10 | 196 | 80 | 753 |
| | 20H3/20K2 | 0 | 0 | 0 | 0 | 0 | 3 | 0 | 0 | 2 | 0 | 0 | 0 | 4 | 0 | 0 | 5 | 2 | 0 | 0 | 0 | 0 | - | 0 | 3 | 2 | 2 | 12 | 3 | 25 |
| | 52H3/52K2 | 0 | 0 | 0 | 0 | 0 | 1 | 0 | 0 | 1 | 0 | 0 | 0 | 1 | 0 | 0 | 1 | 1 | 0 | 0 | 0 | 0 | 0 | 1 | 2 | 1 | 0 | 3 | 1 | 10 |
| | 5E3 | 3 | 1 | 0 | 1 | 2 | 5 | 1 | 0 | 3 | 0 | 0 | 0 | 4 | 0 | 1 | 162 | 54 | 0 | 0 | 0 | 0 | 1 | 21 | 43 | 3 | 3 | 162 | 90 | 746 |
| | 79H2/79K2 | 0 | 0 | 0 | 0 | 0 | 1 | 0 | 0 | 0 | 0 | 0 | 0 | 1 | 0 | 0 | 0 | 0 | 0 | 0 | 0 | 0 | 0 | 0 | 1 | 0 | 0 | 1 | 0 | 2 |
| | 7H3/7K3 | 0 | 0 | 0 | 0 | 0 | 5 | 0 | 0 | 6 | 0 | 0 | 0 | 8 | 0 | 0 | 7 | 2 | 0 | 0 | 0 | 0 | 0 | 2 | 3 | 2 | 2 | 22 | 4 | 107 |
| | 90H3/90K3 | 1 | 0 | 0 | 0 | 0 | 2 | 0 | 0 | 1 | 0 | 0 | 0 | 1 | 0 | 0 | 3 | 1 | 0 | 0 | 0 | 0 | 0 | 1 | 3 | 1 | 1 | 5 | 2 | 17 |
| | A194-01 | 1 | 1 | 0 | 0 | 0 | 2 | 0 | 0 | 1 | 0 | 0 | 0 | 1 | 2 | 13 | 30 | 10 | 0 | 0 | 0 | 0 | 4 | 5 | 10 | 1 | 2 | 37 | 17 | 171 |
| | BJ-03 | 8 | 6 | 0 | 1 | 1 | 21 | 1 | 1 | 26 | 0 | 0 | 3 | 31 | 3 | 35 | 51 | 21 | 0 | 0 | 0 | 0 | 7 | 8 | 24 | 7 | 6 | 56 | 32 | 175 |
| | BJ-76 | 18 | 15 | 0 | 1 | 3 | 77 | 1 | 2 | 60 | 0 | 0 | 4 | 61 | 9 | 84 | 115 | 43 | 0 | 0 | 0 | 0 | 23 | 32 | 128 | 39 | 27 | 227 | 116 | 734 |
| | BTM-1 | 2 | 0 | 0 | 0 | 0 | 5 | 0 | 0 | 6 | 0 | 0 | 1 | 7 | 0 | 0 | 10 | 4 | 0 | 0 | 0 | 0 | 0 | 3 | 7 | 3 | 3 | 16 | 8 | 0 |
| | BTM-8 | 1 | 1 | 0 | 0 | 0 | 3 | 0 | 0 | 4 | 0 | 0 | 0 | 4 | 0 | 0 | 11 | 2 | 0 | 0 | 0 | 0 | 0 | 2 | 4 | 2 | 2 | 13 | 4 | 40 |
| | F-1D7 | 0 | 0 | 0 | 0 | 0 | 0 | 0 | 0 | 0 | 0 | 0 | 0 | 0 | 0 | 0 | 1 | 0 | 0 | 0 | 0 | 0 | 0 | 0 | 1 | 0 | 0 | 1 | 0 | 3 |
| | F-1E7 | 0 | 0 | 0 | 0 | 0 | 0 | 0 | 0 | 0 | 0 | 0 | 0 | 1 | 0 | 1 | 1 | 0 | 0 | 0 | 0 | 0 | 0 | 0 | 1 | 0 | 0 | 2 | 0 | 4 |
| | F-2B4 | 0 | 0 | 0 | 0 | 0 | 0 | 0 | 0 | 0 | 0 | 0 | 0 | 0 | 0 | 0 | 0 | 0 | 0 | 0 | 0 | 0 | 0 | 0 | 0 | 0 | 0 | 0 | 0 | 0 |
| | F-3E2 | 0 | 0 | 0 | 0 | 0 | 0 | 0 | 0 | 0 | 0 | 0 | 0 | 0 | 0 | 0 | 0 | 0 | 0 | 0 | 0 | 0 | 0 | 0 | 0 | 0 | 0 | 0 | 0 | 0 |
| | FDX-01 | 12 | 8 | 0 | 1 | 2 | 25 | 1 | 4 | 32 | 1 | 0 | 6 | 27 | 4 | 77 | 55 | 27 | 0 | 0 | 0 | 0 | 11 | 13 | 33 | 16 | 13 | 69 | 42 | 420 |
| | FIND28 | 3 | 2 | 0 | 0 | 1 | 9 | 0 | 2 | 12 | 0 | 0 | 3 | 21 | 1 | 16 | 20 | 8 | 0 | 0 | 0 | 0 | 2 | 4 | 10 | 7 | 6 | 33 | 12 | 189 |
| | MCD022 Fab | 1 | 1 | 0 | 0 | 0 | 3 | 0 | 0 | 2 | 0 | 0 | 0 | 0 | 0 | 0 | 0 | 0 | 0 | 0 | 0 | 0 | 0 | 0 | 20 | 1 | 2 | 40 | 18 | 0 |
| | MCD022 Fab2 | 2 | 1 | 0 | 0 | 0 | 3 | 0 | 0 | 2 | 0 | 0 | 0 | 0 | 0 | 0 | 0 | 0 | 0 | 0 | 0 | 0 | 0 | 0 | 20 | 1 | 0 | 6 | 9 | 1 |
| | MCD024 Fab | 27 | 17 | 0 | 1 | 2 | 54 | 1 | 3 | 64 | 1 | 0 | 8 | 144 | 0 | 0 | 170 | 56 | 0 | 0 | 0 | 0 | 0 | 18 | 73 | 29 | 25 | 84 | 51 | 446 |
| | MCD024 Fab2 | 21 | 14 | 0 | 1 | 2 | 64 | 1 | 2 | 78 | 0 | 0 | 6 | 156 | 0 | 0 | 125 | 46 | 0 | 0 | 0 | 0 | 0 | 24 | 79 | 31 | 20 | 84 | 20 | 351 |
| | S4-20 | 28 | 8 | 0 | 1 | 3 | 58 | 2 | 3 | 76 | 0 | 0 | 8 | 100 | 17 | 169 | 150 | 51 | 0 | 0 | 0 | 0 | 46 | 56 | 70 | 25 | 31 | 84 | 59 | 603 |

### S11 Table. All Lineages Pooled S - N ratio vs RAP

| | | Detector antibody | | | | | | | | | | | | | | | | | | | | | | | | | | | |
|---|---|---|---|---|---|---|---|---|---|---|---|---|---|---|---|---|---|---|---|---|---|---|---|---|---|---|---|---|---|
| | | 11H2/ 11K1 | 15H3/ 15K3 | 16H2/ 16K1 | 17H2/ 17K3 | 18H2/ 18K2 | 1E7 | 20H3/ 20K2 | 52H3/ 52K2 | 5E3 | 79H2/ 79K2 | 7H3/ 7K3 | 90H3/ 90K3 | A194 -01 | BJ -03 | BJ -76 | BTM -1 | BTM -8 | F-1 D7 | F-1 E7 | F -2B4 | F-3 E2 | FDX -01 | FIND 28 | KI24 | MCD 022 Fab | MCD 022 Fab2 | MCD 024 Fab | MCD 024 Fab2 | S4-20 |
| Capture antibody | 11H2/11K1 | 0 | 0 | 0 | 0 | 0 | 2 | 0 | 1 | 4 | 0 | 0 | 1 | 17 | 0 | 2 | 4 | 1 | 0 | 0 | 0 | 0 | 1 | 1 | 6 | 4 | 3 | 5 | 2 | 16 |
| | 15H3/15K3 | 0 | 0 | 0 | 0 | 0 | 0 | 0 | 0 | 0 | 0 | 0 | 0 | 1 | 0 | 0 | 0 | 0 | 0 | 0 | 0 | 0 | 0 | 0 | 0 | 0 | 0 | 0 | 0 | 0 |
| | 16H2/16K1 | 0 | 0 | 0 | 0 | 0 | 1 | 0 | 0 | 2 | 0 | 0 | 0 | 12 | 0 | 2 | 4 | 1 | 0 | 0 | 0 | 0 | 1 | 0 | 2 | 3 | 2 | 4 | 1 | 15 |
| | 17H2/17K3 | 0 | 0 | 0 | 0 | 0 | 0 | 0 | 0 | 0 | 0 | 0 | 0 | 1 | 0 | 0 | 0 | 0 | 0 | 0 | 0 | 0 | 0 | 0 | 1 | 1 | 0 | 1 | 0 | 0 |
| | 18H2/18K2 | 1 | 0 | 0 | 0 | 0 | 2 | 0 | 0 | 4 | 0 | 0 | 1 | 17 | 0 | 1 | 4 | 1 | 0 | 0 | 0 | 0 | 0 | 1 | 6 | 4 | 3 | 4 | 2 | 15 |
| | 1E7 | 4 | 3 | 0 | 0 | 0 | 9 | 0 | 2 | 13 | 0 | 0 | 4 | 95 | 2 | 12 | 19 | 6 | 0 | 0 | 0 | 0 | 5 | 3 | 18 | 16 | 15 | 17 | 9 | 74 |
| | 20H3/20K2 | 0 | 0 | 0 | 0 | 0 | 0 | 0 | 0 | 1 | 0 | 0 | 0 | 5 | 0 | 0 | 1 | 0 | 0 | 0 | 0 | 0 | - | 0 | 1 | 1 | 1 | 1 | 0 | 4 |
| | 52H3/52K2 | 0 | 0 | 0 | 0 | 0 | 0 | 0 | 0 | 0 | 0 | 0 | 0 | 5 | 0 | 0 | 0 | 0 | 0 | 0 | 0 | 0 | 0 | 0 | 1 | 1 | 0 | 0 | 0 | 1 |
| | 5E3 | 7 | 4 | 0 | 0 | 1 | 12 | 0 | 2 | 22 | 0 | 0 | 4 | 111 | 4 | 16 | 24 | 8 | 0 | 0 | 0 | 0 | 9 | 4 | 28 | 20 | 16 | 20 | 11 | 83 |
| | 79H2/79K2 | 0 | 0 | 0 | 0 | 0 | 0 | 0 | 0 | 0 | 0 | 0 | 0 | 3 | 0 | 0 | 0 | 0 | 0 | 0 | 0 | 0 | 0 | 0 | 0 | 0 | 0 | 0 | 0 | 0 |
| | 7H3/7K3 | 0 | 0 | 0 | 0 | 0 | 1 | 0 | 0 | 1 | 0 | 0 | 0 | 6 | 0 | 1 | 1 | 0 | 0 | 0 | 0 | 0 | 0 | 0 | 1 | 1 | 1 | 2 | 0 | 11 |
| | 90H3/90K3 | 0 | 0 | 0 | 0 | 0 | 1 | 0 | 0 | 1 | 0 | 0 | 0 | 8 | 0 | 0 | 0 | 0 | 0 | 0 | 0 | 0 | 0 | 0 | 1 | 1 | 1 | 1 | 1 | 2 |
| | A194-01 | 2 | 1 | 0 | 0 | 0 | 5 | 0 | 1 | 5 | 0 | 0 | 3 | 52 | 0 | 2 | 5 | 2 | 0 | 0 | 0 | 0 | 1 | 2 | 12 | 7 | 7 | 6 | 3 | 20 |
| | BJ-03 | 3 | 2 | 0 | 0 | 0 | 5 | 0 | 1 | 11 | 0 | 0 | 1 | 43 | 2 | 7 | 11 | 4 | 0 | 0 | 0 | 0 | 2 | 2 | 5 | 6 | 5 | 9 | 5 | 43 |
| | BJ-76 | 3 | 2 | 0 | 0 | 0 | 9 | 0 | 0 | 14 | 0 | 0 | 1 | 47 | 2 | 9 | 14 | 5 | 0 | 0 | 0 | 0 | 3 | 4 | 12 | 9 | 6 | 16 | 8 | 71 |
| | BTM-1 | 0 | 0 | 0 | 0 | 0 | 1 | 0 | 0 | 1 | 0 | 0 | 0 | 4 | 0 | 0 | 1 | 0 | 0 | 0 | 0 | 0 | 0 | 0 | 1 | 1 | 1 | 2 | 1 | 1 |
| | BTM-8 | 0 | 0 | 0 | 0 | 0 | 0 | 0 | 0 | 1 | 0 | 0 | 0 | 2 | 0 | 0 | 1 | 0 | 0 | 0 | 0 | 0 | 0 | 0 | 1 | 0 | 0 | 1 | 0 | 4 |
| | F-1D7 | 0 | 0 | 0 | 0 | 0 | 0 | 0 | 0 | 0 | 0 | 0 | 0 | 0 | 0 | 0 | 0 | 0 | 0 | 0 | 0 | 0 | 0 | 0 | 0 | 0 | 0 | 0 | 0 | 0 |
| | F-1E7 | 0 | 0 | 0 | 0 | 0 | 0 | 0 | 0 | 0 | 0 | 0 | 0 | 0 | 0 | 0 | 0 | 0 | 0 | 0 | 0 | 0 | 0 | 0 | 0 | 0 | 0 | 0 | 0 | 0 |
| | F-2B4 | 0 | 0 | 0 | 0 | 0 | 0 | 0 | 0 | 0 | 0 | 0 | 0 | 0 | 0 | 0 | 0 | 0 | 0 | 0 | -5 | 0 | 0 | 0 | 0 | 0 | 0 | 0 | 0 | 0 |
| | F-3E2 | 0 | 0 | 0 | 0 | 0 | 0 | 0 | 0 | 0 | 0 | 0 | 0 | 0 | 0 | 0 | 0 | 0 | 0 | 0 | 0 | 0 | 0 | 0 | 0 | 0 | 0 | 0 | 0 | 0 |
| | FDX-01 | 2 | 1 | 0 | 0 | 0 | 7 | 0 | 1 | 9 | 0 | 0 | 3 | 69 | 1 | 7 | 7 | 3 | 0 | 0 | 0 | 0 | 1 | 3 | 12 | 10 | 6 | 10 | 4 | 41 |
| | FIND28 | 1 | 0 | 0 | 0 | 0 | 2 | 0 | 0 | 2 | 0 | 0 | 1 | 20 | 0 | 2 | 2 | 1 | 0 | 0 | 0 | 0 | 0 | 1 | 4 | 4 | 2 | 4 | 1 | 11 |
| | MCD022 Fab | 2 | 1 | 0 | 0 | 0 | 8 | 0 | 2 | 6 | 0 | 0 | 4 | 50 | 0 | 1 | 1 | 0 | 0 | 0 | 0 | 0 | 0 | 1 | 16 | 9 | 9 | 7 | 3 | 6 |
| | MCD022 Fab2 | 2 | 1 | 0 | 0 | 0 | 6 | 0 | 1 | 7 | 0 | 0 | 2 | 45 | 0 | 1 | 2 | 1 | 0 | 0 | 0 | 0 | 1 | 1 | 12 | 7 | 8 | 9 | 4 | 12 |
| | MCD024 Fab | 4 | 3 | 0 | 0 | 0 | 12 | 0 | 1 | 15 | 0 | 0 | 2 | 66 | 1 | 1 | 16 | 6 | 0 | 0 | 0 | 0 | 2 | 5 | 25 | 12 | 10 | 13 | 7 | 55 |
| | MCD024 Fab2 | 4 | 2 | 0 | 0 | 0 | 9 | 0 | 1 | 13 | 0 | 0 | 2 | 61 | 1 | 1 | 14 | 5 | 0 | 0 | 0 | 0 | 1 | 4 | 21 | 11 | 10 | 11 | 6 | 51 |
| | S4-20 | 5 | 2 | 0 | 0 | 0 | 10 | 0 | 1 | 12 | 0 | 0 | 2 | 100 | 4 | 15 | 19 | 7 | 0 | 0 | 0 | 0 | 6 | 8 | 21 | 11 | 10 | 11 | 6 | 79 |

### S12 Table. All Lineages Pooled S/N vs RAP

| Capture antibody | Detector Antibody | | | | | | | | | | | | | | | | | | | | | | | | | | | | |
|---|---|---|---|---|---|---|---|---|---|---|---|---|---|---|---|---|---|---|---|---|---|---|---|---|---|---|---|---|---|
| | 11H2/ 11K1 | 15H3/ 15K3 | 16H2/ 16K1 | 17H2/ 17K3 | 18H2/ 18K2 | 1E7 | 20H3/ 20K2 | 52H3/ 52K2 | 5E3 | 79H2/ 79K2 | 7H3/ 7K3 | 90H3/ 90K3 | A194 -01 | BJ -03 | BJ -76 | BTM -1 | BTM -8 | F-1 D7 | F-1 E7 | F -2B4 | F-3 E2 | FDX -01 | FIND 28 | KI24 | MCD 022 Fab | MCD 022 Fab2 | MCD 024 Fab | MCD 024 Fab2 | S4-20 |
| 11H2/11K1 | | | | | | | | | | | | | | | | | | | | | | | | | | | | | |
| 15H3/15K3 | 1 | 0 | 0 | 0 | 0 | 7 | 0 | 1 | 6 | 0 | 0 | 1 | 6 | 1 | 5 | 12 | 5 | 0 | 0 | 0 | 0 | 5 | 4 | 13 | 6 | 6 | 18 | 8 | 51 |
| 16H2/16K1 | 0 | 0 | 0 | 0 | 0 | 0 | 0 | 0 | 1 | 0 | 0 | 0 | 1 | 0 | 0 | 1 | 0 | 0 | 0 | 0 | 0 | 0 | 0 | 1 | 1 | 0 | 1 | 0 | 2 |
| 17H2/17K3 | 1 | 0 | 0 | 0 | 0 | 3 | 0 | 1 | 8 | 0 | 0 | 2 | 15 | 1 | 4 | 10 | 3 | 0 | 0 | 0 | 0 | 3 | 2 | 7 | 8 | 8 | 16 | 6 | 46 |
| 18H2/18K2 | 0 | 0 | 0 | 0 | 0 | 1 | 0 | 0 | 1 | 0 | 0 | 0 | 1 | 0 | 0 | 0 | 0 | 0 | 0 | 0 | 0 | 0 | 0 | 2 | 2 | 1 | 3 | 1 | 1 |
| 1E7 | 1 | 0 | 0 | 0 | 0 | 8 | 0 | 1 | 8 | 0 | 0 | 2 | 3 | 1 | 1 | 10 | 4 | 0 | 0 | 0 | 0 | 1 | 3 | 16 | 5 | 6 | 14 | 3 | 47 |
| 20H3/20K2 | 5 | 2 | 0 | 1 | 1 | 11 | 1 | 1 | 8 | 0 | 0 | 1 | 13 | 5 | 15 | 53 | 20 | 0 | 1 | 0 | 0 | 16 | 9 | 41 | 11 | 16 | 69 | 36 | 225 |
| 52H3/52K2 | 0 | 0 | 0 | 0 | 0 | 2 | 0 | 0 | 2 | 0 | 0 | 1 | 8 | 0 | 0 | 3 | 1 | 0 | 0 | 0 | 0 | - | 0 | 4 | 2 | 3 | 6 | 1 | 12 |
| 5E3 | 0 | 0 | 0 | 0 | 0 | 1 | 0 | 0 | 1 | 0 | 0 | 1 | 2 | 0 | 0 | 1 | 0 | 0 | 0 | 0 | 0 | 0 | 1 | 3 | 1 | 1 | 2 | 1 | 4 |
| 79H2/79K2 | 3 | 1 | 0 | 1 | 1 | 5 | 1 | 1 | 4 | 0 | 0 | 0 | 5 | 11 | 26 | 66 | 26 | 1 | 2 | 0 | 0 | 24 | 14 | 43 | 6 | 5 | 74 | 45 | 276 |
| 7H3/7K3 | 0 | 0 | 0 | 0 | 0 | 1 | 0 | 0 | 0 | 0 | 0 | 0 | 1 | 0 | 0 | 0 | 0 | 0 | 0 | 0 | 0 | 0 | 0 | 1 | 1 | 0 | 1 | 1 | 2 |
| 90H3/90K3 | 1 | 0 | 0 | 0 | 0 | 3 | 0 | 0 | 3 | 0 | 0 | 0 | 8 | 0 | 2 | 2 | 1 | 0 | 0 | 0 | 0 | 2 | 1 | 2 | 3 | 3 | 7 | 1 | 38 |
| A194-01 | 1 | 1 | 0 | 0 | 0 | 2 | 0 | 0 | 1 | 0 | 0 | 1 | 3 | 0 | 0 | 2 | 1 | 0 | 0 | 0 | 0 | 0 | 1 | 5 | 2 | 2 | 3 | 1 | 8 |
| BJ-03 | 1 | 1 | 0 | 0 | 0 | 3 | 0 | 1 | 2 | 0 | 0 | 0 | 3 | 2 | 7 | 17 | 6 | 0 | 0 | 0 | 0 | 4 | 5 | 17 | 2 | 4 | 23 | 10 | 69 |
| BJ-76 | 9 | 6 | 0 | 1 | 1 | 19 | 1 | 2 | 33 | 0 | 0 | 4 | 44 | 4 | 19 | 29 | 12 | 0 | 0 | 0 | 0 | 6 | 5 | 15 | 15 | 14 | 34 | 20 | 136 |
| BTM-1 | 10 | 9 | 0 | 1 | 1 | 36 | 1 | 2 | 36 | 0 | 0 | 3 | 45 | 5 | 25 | 39 | 15 | 0 | 0 | 0 | 0 | 13 | 11 | 43 | 28 | 21 | 74 | 33 | 267 |
| BTM-8 | 2 | 1 | 0 | 0 | 0 | 3 | 0 | 0 | 4 | 0 | 0 | 1 | 7 | 0 | 1 | 4 | 1 | 0 | 0 | 0 | 0 | 1 | 1 | 5 | 3 | 3 | 6 | 3 | 3 |
| F-1D7 | 1 | 0 | 0 | 0 | 0 | 1 | 0 | 0 | 2 | 0 | 0 | 0 | 4 | 0 | 1 | 3 | 1 | 0 | 0 | 0 | 0 | 1 | 1 | 3 | 2 | 2 | 4 | 2 | 13 |
| F-1E7 | 0 | 0 | 0 | 0 | 0 | 0 | 0 | 0 | 0 | 0 | 0 | 0 | 0 | 0 | 0 | 0 | 0 | 0 | 0 | 0 | 0 | 0 | 0 | 0 | 0 | 0 | 1 | 0 | 1 |
| F-2B4 | 0 | 0 | 0 | 0 | 0 | 0 | 0 | 0 | 0 | 0 | 0 | 0 | 0 | 0 | 0 | 0 | 0 | 0 | 0 | 0 | 0 | 0 | 0 | 0 | 0 | 0 | 1 | 0 | 1 |
| F-3E2 | 0 | 0 | 0 | 0 | 0 | 0 | 0 | 0 | 0 | 0 | 0 | 0 | 0 | 0 | 0 | 0 | 0 | 0 | 0 | 0 | 0 | 0 | 0 | 0 | 0 | 0 | 0 | 0 | 0 |
| FDX-01 | 0 | 0 | 0 | 0 | 0 | 0 | 0 | 0 | 0 | 0 | 0 | 0 | 0 | 0 | 0 | 0 | 0 | 0 | 0 | 0 | 0 | 0 | 0 | 0 | 0 | 0 | 0 | 0 | 0 |
| FIND28 | 7 | 5 | 0 | 1 | 1 | 23 | 0 | 4 | 28 | 1 | 0 | 8 | 41 | 3 | 24 | 23 | 10 | 0 | 0 | 0 | 0 | 6 | 9 | 27 | 24 | 18 | 39 | 20 | 177 |
| MCD022 Fab | 3 | 2 | 0 | 0 | 1 | 7 | 0 | 2 | 9 | 0 | 0 | 4 | 26 | 1 | 7 | 9 | 3 | 0 | 0 | 0 | 0 | 2 | 2 | 8 | 12 | 7 | 16 | 5 | 51 |
| MCD022 Fab2 | 2 | 1 | 0 | 0 | 0 | 4 | 0 | 1 | 2 | 0 | 0 | 1 | 4 | 1 | 2 | 3 | 1 | 0 | 0 | 0 | 0 | 2 | 3 | 33 | 3 | 5 | 26 | 5 | 5 |
| MCD024 Fab | 2 | 1 | 0 | 0 | 0 | 4 | 0 | 1 | 2 | 0 | 0 | 0 | 4 | 1 | 2 | 4 | 2 | 0 | 0 | 0 | 0 | 3 | 1 | 30 | 2 | 1 | 2 | 3 | 7 |
| MCD024 Fab2 | 18 | 11 | 0 | 1 | 2 | 45 | 1 | 3 | 51 | 1 | 0 | 8 | 175 | 5 | 5 | 64 | 26 | 0 | 0 | 0 | 0 | 5 | 19 | 86 | 22 | 36 | 41 | 10 | 192 |
| S4-20 | 14 | 8 | 0 | 1 | 2 | 46 | 1 | 3 | 57 | 0 | 0 | 6 | 166 | 3 | 4 | 49 | 18 | 0 | 0 | 0 | 0 | 4 | 16 | 70 | 30 | 23 | 34 | 6 | 181 |
| | 16 | 8 | 0 | 1 | 2 | 35 | 1 | 3 | 46 | 0 | 0 | 6 | 100 | 10 | 42 | 51 | 20 | 0 | 0 | 0 | 0 | 26 | 28 | 65 | 25 | 31 | 34 | 23 | 226 |

### S13 Table. uLAM S-N vs RAP

| | Detector Antibody | | | | | | | | | | | | | | | | | | | | | | | | | | | | |
|---|---|---|---|---|---|---|---|---|---|---|---|---|---|---|---|---|---|---|---|---|---|---|---|---|---|---|---|---|---|
| Capture antibody | 11H2/ 11K1 | 15H3/ 15K3 | 16H2/ 16K1 | 17H2/ 17K3 | 18H2/ 18K2 | 1E7 | 20H3/ 20K2 | 52H3/ 52K2 | 5E3 | 79H2/ 79K2 | 7H3/ 7K3 | 90H3/ 90K3 | A194 -01 | BJ -03 | BJ -76 | BTM -1 | BTM -8 | F-1 D7 | F-1 E7 | F -2B4 | F-3 E2 | FDX -01 | FIND 28 | KI24 | MCD 022 Fab | MCD 022 Fab2 | MCD 024 Fab | MCD 024 Fab2 | S4-20 |
| 11H2/11K1 | 0 | 0 | 0 | 0 | 0 | 60 | 0 | 33 | 22 | 5 | 0 | 54 | 607 | 0 | 1 | 4 | 2 | 0 | 0 | -1 | 0 | 4 | 12 | 57 | 79 | 41 | 2 | 1 | 7 |
| 15H3/15K3 | 0 | 0 | 0 | 0 | 0 | 1 | 0 | 1 | 0 | 0 | 0 | 1 | 12 | 0 | 0 | 0 | 0 | 0 | 0 | -1 | 0 | 0 | 0 | 2 | 2 | 1 | 0 | 0 | 0 |
| 16H2/16K1 | 0 | 1 | 0 | 0 | 0 | 8 | 0 | 14 | 11 | 7 | 0 | 24 | 381 | 0 | 0 | 0 | 0 | 0 | 0 | -5 | 0 | 0 | 2 | 8 | 48 | 12 | 0 | 0 | 0 |
| 17H2/17K3 | 0 | 0 | 0 | 0 | 0 | 1 | 0 | 1 | 1 | 0 | 0 | 1 | 3 | 0 | 0 | 0 | 0 | 0 | 0 | 0 | 0 | 0 | 0 | 0 | 1 | 0 | 0 | 0 | 0 |
| 18H2/18K2 | 0 | 0 | 0 | 0 | 0 | 48 | 0 | 17 | 9 | 4 | 0 | 30 | 410 | 0 | 0 | 2 | 1 | 0 | 0 | -2 | 0 | 3 | 4 | 24 | 46 | 28 | 1 | 0 | 3 |
| 1E7 | 12 | 25 | 0 | 1 | 5 | 77 | 1 | 14 | 4 | 3 | 0 | 52 | 1111 | 0 | 5 | 9 | 4 | 0 | 0 | -2 | 0 | 8 | 6 | 30 | 98 | 105 | 5 | 4 | 27 |
| 20H3/20K2 | 0 | 0 | 0 | 0 | 0 | 3 | 0 | 3 | 3 | 0 | 0 | 5 | 32 | 0 | 0 | 0 | 0 | 0 | 0 | -1 | 0 | - | 0 | 2 | 5 | 1 | 0 | 0 | 0 |
| 52H3/52K2 | 1 | 1 | 0 | 0 | 0 | 26 | 0 | 9 | 3 | 2 | 0 | 18 | 240 | 0 | 0 | 1 | 0 | 0 | 0 | -2 | 0 | 1 | 4 | 19 | 25 | 16 | 0 | 0 | 1 |
| 5E3 | 205 | 167 | 1 | 19 | 55 | 196 | 21 | 64 | 39 | 6 | 4 | 98 | 2745 | 0 | 5 | 18 | 8 | 0 | 0 | -2 | 0 | 17 | 44 | 222 | 305 | 165 | 9 | 7 | 40 |
| 79H2/79K2 | 0 | 0 | 0 | 0 | 0 | 11 | 0 | 3 | 1 | 1 | 0 | 7 | 87 | 0 | 0 | 0 | 0 | 0 | 0 | -1 | 0 | 0 | 1 | 4 | 10 | 6 | 0 | 0 | 0 |
| 7H3/7K3 | 0 | 0 | 0 | 0 | 0 | 3 | 0 | 2 | 1 | 0 | 0 | 3 | 22 | 0 | 0 | 0 | 0 | 0 | 0 | -2 | 0 | 0 | 1 | 6 | 3 | 2 | 0 | 0 | 0 |
| 90H3/90K3 | 1 | 1 | 0 | 0 | 0 | 18 | 0 | 5 | 2 | 1 | 0 | 11 | 163 | 0 | 0 | 1 | 0 | 0 | 0 | -1 | 0 | 1 | 4 | 18 | 14 | 12 | 0 | 0 | 1 |
| A194-01 | 16 | 24 | 0 | 1 | 4 | 106 | 1 | 28 | 9 | 5 | 0 | 58 | 1137 | 0 | 2 | 18 | 7 | 0 | 0 | 0 | 0 | 13 | 25 | 132 | 103 | 105 | 7 | 4 | 26 |
| BJ-03 | 0 | 0 | 0 | 0 | 0 | 0 | 0 | 0 | 0 | 0 | 0 | 0 | 0 | 0 | 0 | 0 | 0 | 0 | 0 | -6 | 0 | 0 | 1 | 0 | 0 | 0 | 0 | 0 | 0 |
| BJ-76 | 0 | 1 | 0 | 0 | 0 | 1 | 0 | 0 | 1 | 0 | 0 | 1 | 9 | 0 | 0 | 2 | 1 | 0 | 0 | -1 | 0 | 0 | 0 | 0 | 1 | 1 | 1 | 1 | 3 |
| BTM-1 | 0 | 0 | 0 | 0 | 0 | 0 | 0 | 0 | 0 | 0 | 0 | 0 | 8 | 0 | 0 | 0 | 0 | 0 | 0 | -1 | 0 | 0 | 0 | 0 | 1 | 1 | 0 | 0 | 1 |
| BTM-8 | 0 | 0 | 0 | 0 | 0 | 0 | 0 | 0 | 0 | 0 | 0 | 0 | 4 | 0 | 0 | 0 | 0 | 0 | 0 | -3 | 0 | 0 | 0 | 1 | 0 | 0 | 0 | 0 | 0 |
| F-1D7 | 0 | 0 | 0 | 0 | 0 | 0 | 0 | 0 | 0 | 0 | 0 | 0 | 0 | 0 | 0 | 0 | 0 | 0 | 0 | -5 | 0 | 0 | 0 | 0 | 0 | 0 | 0 | 0 | 0 |
| F-1E7 | 0 | 0 | 0 | 0 | 0 | 0 | 0 | 0 | 0 | 0 | 0 | 0 | 2 | 0 | 0 | 0 | 0 | 0 | 0 | 2 | 0 | 0 | 0 | 0 | 0 | 0 | 0 | 0 | 0 |
| F-2B4 | 0 | 0 | 0 | 0 | 0 | 0 | 0 | 0 | 0 | 0 | 0 | 0 | 39 | 0 | 0 | 0 | 0 | 0 | 0 | -40 | 0 | 0 | 0 | 0 | 0 | 0 | 0 | 0 | 0 |
| F-3E2 | 0 | 0 | 0 | 0 | 0 | 0 | 0 | 0 | 0 | 0 | 0 | 0 | 1 | 0 | 0 | 0 | 0 | 0 | 0 | 1 | 0 | 0 | 0 | 0 | 0 | 0 | 0 | 0 | 0 |
| FDX-01 | 7 | 21 | 0 | 0 | 3 | 177 | 0 | 59 | 22 | 13 | 0 | 150 | 2067 | 0 | 1 | 1 | 0 | 0 | 0 | -5 | 0 | 9 | 51 | 175 | 210 | 119 | 0 | 0 | 1 |
| FIND28 | 7 | 8 | 0 | 0 | 1 | 55 | 0 | 26 | 7 | 4 | 0 | 56 | 661 | 0 | 0 | 0 | 0 | 0 | 0 | -2 | 0 | 3 | 35 | 85 | 83 | 25 | 0 | 0 | 0 |
| MCD022 Fab | 10 | 21 | 0 | 1 | 4 | 206 | 1 | 61 | 17 | 10 | 0 | 114 | 1911 | 0 | 1 | 10 | 4 | 0 | 0 | -1 | 0 | 13 | 53 | 206 | 182 | 163 | 6 | 4 | 12 |
| MCD022 Fab2 | 10 | 31 | 0 | 0 | 3 | 32 | 1 | 5 | 1 | 1 | 0 | 10 | 961 | 0 | 2 | 19 | 8 | 0 | 0 | -1 | 0 | 5 | 12 | 14 | 26 | 35 | 11 | 8 | 26 |
| MCD024 Fab | 1 | 0 | 0 | 0 | 0 | 1 | 0 | 1 | 2 | 0 | 0 | 3 | 53 | 0 | 0 | 2 | 1 | 0 | 0 | -1 | 0 | 0 | 0 | 1 | 5 | 4 | 1 | 0 | 2 |
| MCD024 Fab2 | 1 | 0 | 0 | 0 | 0 | 1 | 0 | 1 | 1 | 0 | 0 | 3 | 51 | 0 | 0 | 0 | 0 | 0 | 0 | 0 | 0 | 0 | 0 | 1 | 5 | 5 | 0 | 0 | 1 |
| S4-20 | 1 | 1 | 0 | 0 | 0 | 2 | 0 | 1 | 2 | 0 | 0 | 4 | 100 | 0 | 0 | 2 | 1 | 0 | 0 | -2 | 0 | 0 | 1 | 2 | 7 | 6 | 0 | 0 | 2 |

### S14 Table. uLAM S/N vs RAP

| | Detector Antibody | | | | | | | | | | | | | | | | | | | | | | | | | | | | |
|---|---|---|---|---|---|---|---|---|---|---|---|---|---|---|---|---|---|---|---|---|---|---|---|---|---|---|---|---|---|
| Capture antibody | 11H2/ 11K1 | 15H3/ 15K3 | 16H2/ 16K1 | 17H2/ 17K3 | 18H2/ 18K2 | 1E7 | 20H3/ 20K2 | 52H3/ 52K2 | 5E3 | 79H2/ 79K2 | 7H3/ 7K3 | 90H3/ 90K3 | A194 -01 | BJ -03 | BJ -76 | BTM -1 | BTM -8 | F-1 D7 | F-1 E7 | F -2B4 | F-3 E2 | FDX -01 | FIND 28 | KI24 | MCD 022 Fab | MCD 022 Fab2 | MCD 024 Fab | MCD 024 Fab2 | S4-20 |
| 11H2/11K1 | 2 | 2 | 2 | 2 | 2 | 122 | 2 | 43 | 21 | 11 | 2 | 34 | 130 | 2 | 3 | 13 | 6 | 1 | 2 | 1 | 1 | 14 | 33 | 94 | 65 | 57 | 5 | 5 | 19 |
| 15H3/15K3 | 1 | 2 | 1 | 1 | 2 | 7 | 2 | 4 | 2 | 2 | 2 | 4 | 15 | 2 | 2 | 1 | 2 | 2 | 2 | 1 | 2 | 2 | 3 | 7 | 5 | 4 | 2 | 2 | 2 |
| 16H2/16K1 | 2 | 4 | 2 | 2 | 2 | 25 | 2 | 48 | 27 | 21 | 2 | 52 | 314 | 2 | 2 | 2 | 3 | 2 | 1 | 1 | 2 | 2 | 7 | 23 | 108 | 34 | 2 | 3 | 3 |
| 17H2/17K3 | 2 | 2 | 2 | 2 | 2 | 4 | 2 | 5 | 2 | 3 | 1 | 6 | 4 | 1 | 2 | 2 | 2 | 1 | 1 | 2 | 1 | 1 | 1 | 3 | 5 | 2 | 2 | 1 | 2 |
| 18H2/18K2 | 2 | 2 | 2 | 2 | 2 | 103 | 2 | 34 | 13 | 9 | 2 | 33 | 137 | 2 | 3 | 7 | 4 | 2 | 2 | 1 | 2 | 11 | 13 | 62 | 50 | 46 | 4 | 3 | 10 |
| 1E7 | 13 | 17 | 2 | 5 | 12 | 73 | 5 | 7 | 3 | 3 | 2 | 7 | 112 | 2 | 12 | 18 | 11 | 2 | 2 | 1 | 1 | 30 | 19 | 66 | 42 | 111 | 16 | 14 | 63 |
| 20H3/20K2 | 1 | 2 | 2 | 2 | 2 | 14 | 1 | 11 | 6 | 3 | 1 | 16 | 43 | 2 | 1 | 2 | 2 | 1 | 2 | 1 | 1 | - | 2 | 7 | 12 | 5 | 2 | 1 | 3 |
| 52H3/52K2 | 4 | 5 | 1 | 2 | 2 | 56 | 2 | 15 | 6 | 5 | 2 | 18 | 94 | 2 | 2 | 3 | 3 | 2 | 2 | 1 | 2 | 7 | 11 | 55 | 34 | 29 | 2 | 2 | 4 |
| 5E3 | 48 | 33 | 4 | 39 | 64 | 43 | 38 | 12 | 6 | 4 | 12 | 6 | 82 | 2 | 14 | 37 | 22 | 2 | 2 | 1 | 2 | 45 | 135 | 304 | 42 | 51 | 28 | 25 | 102 |
| 79H2/79K2 | 3 | 3 | 2 | 2 | 2 | 39 | 2 | 9 | 2 | 3 | 2 | 12 | 61 | 2 | 2 | 3 | 3 | 1 | 2 | 1 | 2 | 3 | 4 | 16 | 21 | 16 | 2 | 2 | 2 |
| 7H3/7K3 | 2 | 2 | 2 | 2 | 2 | 12 | 2 | 8 | 4 | 3 | 2 | 8 | 29 | 2 | 2 | 2 | 2 | 2 | 2 | 1 | 1 | 2 | 5 | 17 | 9 | 7 | 2 | 2 | 2 |
| 90H3/90K3 | 3 | 3 | 1 | 2 | 2 | 41 | 2 | 12 | 4 | 4 | 2 | 13 | 68 | 2 | 2 | 4 | 2 | 1 | 1 | 1 | 1 | 5 | 14 | 61 | 23 | 24 | 2 | 2 | 6 |
| A194-01 | 12 | 10 | 2 | 4 | 6 | 47 | 4 | 8 | 4 | 4 | 2 | 6 | 44 | 2 | 10 | 38 | 23 | 2 | 3 | 2 | 3 | 56 | 64 | 161 | 20 | 50 | 19 | 15 | 65 |
| BJ-03 | 2 | 2 | 2 | 2 | 2 | 2 | 2 | 2 | 2 | 2 | 2 | 2 | 2 | 1 | 2 | 2 | 2 | 2 | 2 | 1 | 2 | 2 | 4 | 2 | 2 | 2 | 2 | 2 | 2 |
| BJ-76 | 3 | 4 | 2 | 2 | 2 | 5 | 2 | 2 | 4 | 2 | 2 | 4 | 9 | 1 | 2 | 6 | 4 | 2 | 2 | 1 | 2 | 2 | 2 | 3 | 3 | 4 | 4 | 4 | 10 |
| BTM-1 | 3 | 3 | 2 | 2 | 2 | 3 | 2 | 2 | 3 | 2 | 2 | 3 | 16 | 2 | 2 | 2 | 2 | 2 | 1 | 1 | 2 | 2 | 3 | 3 | 4 | 5 | 3 | 3 | 3 |
| BTM-8 | 2 | 2 | 2 | 2 | 1 | 3 | 2 | 2 | 2 | 2 | 1 | 2 | 8 | 1 | 2 | 2 | 2 | 1 | 1 | 1 | 2 | 2 | 3 | 4 | 3 | 3 | 1 | 2 | 3 |
| F-1D7 | 2 | 2 | 2 | 2 | 2 | 2 | 2 | 3 | 2 | 2 | 2 | 2 | 2 | 1 | 2 | 2 | 1 | 2 | 2 | 1 | 2 | 2 | 2 | 2 | 2 | 2 | 2 | 2 | 2 |
| F-1E7 | 2 | 2 | 2 | 2 | 2 | 2 | 2 | 2 | 2 | 2 | 2 | 2 | 3 | 2 | 2 | 2 | 2 | 2 | 2 | 2 | 2 | 2 | 2 | 2 | 2 | 2 | 2 | 2 | 2 |
| F-2B4 | 2 | 2 | 2 | 2 | 2 | 2 | 2 | 2 | 2 | 2 | 2 | 2 | 26 | 2 | 2 | 2 | 2 | 2 | 2 | 2 | 2 | 2 | 2 | 2 | 2 | 2 | 2 | 2 | 2 |
| F-3E2 | 2 | 2 | 1 | 2 | 1 | 1 | 1 | 1 | 1 | 1 | 1 | 2 | 2 | 1 | 1 | 1 | 1 | 1 | 1 | 3 | 1 | 1 | 1 | 2 | 2 | 1 | 1 | 2 | 1 |
| FDX-01 | 22 | 70 | 2 | 3 | 13 | 455 | 3 | 194 | 65 | 44 | 2 | 399 | 1181 | 1 | 4 | 5 | 3 | 2 | 1 | 1 | 2 | 27 | 132 | 475 | 396 | 314 | 3 | 3 | 5 |
| FIND28 | 27 | 31 | 2 | 3 | 6 | 200 | 2 | 90 | 22 | 13 | 2 | 191 | 519 | 2 | 3 | 3 | 2 | 2 | 2 | 1 | 2 | 13 | 78 | 146 | 191 | 74 | 2 | 2 | 3 |
| MCD022 Fab | 12 | 13 | 2 | 3 | 10 | 106 | 3 | 19 | 7 | 8 | 3 | 14 | 154 | 2 | 5 | 31 | 17 | 2 | 1 | 1 | 2 | 44 | 169 | 477 | 46 | 86 | 19 | 13 | 36 |
| MCD022 Fab2 | 12 | 16 | 1 | 2 | 6 | 22 | 2 | 4 | 2 | 2 | 3 | 3 | 67 | 2 | 5 | 28 | 21 | 2 | 2 | 1 | 2 | 18 | 9 | 40 | 5 | 3 | 2 | 6 | 76 |
| MCD024 Fab | 4 | 3 | 2 | 2 | 3 | 6 | 2 | 5 | 5 | 2 | 2 | 14 | 130 | 2 | 1 | 9 | 4 | 1 | 2 | 1 | 2 | 2 | 3 | 5 | 18 | 16 | 3 | 3 | 9 |
| MCD024 Fab2 | 4 | 3 | 2 | 2 | 2 | 6 | 2 | 4 | 6 | 2 | 2 | 10 | 116 | 2 | 2 | 3 | 3 | 1 | 2 | 2 | 2 | 2 | 3 | 5 | 14 | 11 | 2 | 2 | 4 |
| S4-20 | 6 | 4 | 2 | 2 | 2 | 8 | 2 | 6 | 7 | 3 | 2 | 12 | 100 | 2 | 2 | 5 | 3 | 2 | 1 | 1 | 2 | 2 | 3 | 6 | 17 | 20 | 2 | 2 | 5 |
